# Asynchronous origins of Yellow-Stripe transporters and Nicotianamine Synthase underlie the evolution of plant Strategy-II iron uptake

**DOI:** 10.64898/2026.08.27.747422

**Authors:** Matheus L. C. B. de Campos, Wenderson F. C. Rodrigues, Helena F. Gomes, Felipe K. Ricachenevsky, Joni E. Lima, Luiz-Eduardo Del-Bem

## Abstract

Plants acquire iron using two canonical mechanisms: reduction-based uptake (Strategy I) and chelation-based uptake (Strategy II). Although Strategy II is characteristic of grasses, genes associated with chelated-metal transport occur broadly across plant lineages, obscuring how this pathway evolved. Here, we reconstruct the evolutionary history of Yellow Stripe-Like (YSL) transporters and Nicotianamine Synthase (NAS) across Archaeplastida and their closest non-plant homologs. Phylogenomic, distributional, and structural analyses reveal pronounced temporal uncoupling between these gene families. YSL most likely originated early in Viridiplantae and represents the deepest evolutionary module of the chelated-metal transport system. In contrast, NAS appeared much later through independent horizontal gene transfer events from fungi into euphyllophytes and specific moss lineages. This timing indicates that YSL-mediated transport initially functioned independently of nicotianamine, implying ancestral use of alternative siderophores. A pronounced expansion of YSL genes in Poaceae coincides with the emergence of canonical Strategy II, suggesting that this pathway arose by exaptation of pre-existing chelated-metal transport mechanisms.

## Introduction

Iron (Fe) has been essential to life since the emergence of early metabolic systems on Earth (Keller et al., 2014; Aulakh et al., 2022). Its ability to alternate between ferrous (Fe²⁺) and ferric (Fe³⁺) states provides a broad range of redox potentials that support respiration, photosynthesis, nitrogen fixation, ribonucleotide reduction, and other central reactions (Guerinot and Yi, 1994). The same redox chemistry that makes Fe useful also makes it potentially toxic: Fe²⁺/Fe³⁺ cycling drives Fenton chemistry and generates highly reactive hydroxyl radicals, requiring strict regulation of uptake, intracellular trafficking, and storage (Guerinot and Yi, 1994; Eide et al., 1996). For plants, this biochemical demand is compounded by a geochemical constraint. In neutral-to-alkaline aerobic soils, Fe occurs predominantly as Fe³⁺ and forms poorly soluble oxyhydroxides, whereas Fe²⁺ is more soluble but unstable under oxic conditions and is mainly available in reducing or low-oxygen environments (Guerinot and Yi, 1994; Hoffmann et al., 2012; Kobayashi and Nishizawa, 2012; Mimmo et al., 2014; Connorton et al., 2017). Plants therefore require mechanisms that mobilize, transport, and safely store this scarce micronutrient while limiting metal-induced oxidative damage.

Land plants (Embryophyta) use two canonical Fe-uptake strategies (Marschner et al., 1986; Römheld and Marschner, 1986; Huang et al., 2024). Strategy I, used by most nongrass plants, relies on rhizosphere acidification and plasma-membrane ferric reduction by Ferric Reductase Oxidase (FRO2), followed by Fe²⁺ uptake by Iron-Regulated Transporter (IRT) proteins (Eide et al., 1996; Vert et al., 2002). Strategy II, characteristic of grasses (Poaceae), mobilizes Fe³⁺ through secretion of mugineic acid family phytosiderophores (PS). A complete Strategy II response requires coordinated synthesis of nicotianamine (NA) by Nicotianamine Synthase (NAS) (Higuchi et al., 1994; Higuchi et al., 1999), conversion of NA into deoxymugineic acid (DMA) by Nicotianamine Aminotransferase (NAAT) and Deoxymugineic Acid Synthase (DMAS), export of DMA to the rhizosphere by TOM transporters such as TOM1 and TOM2, and uptake of Fe³⁺-PS complexes by Yellow Stripe-Like (YSL) transporters (Takagi et al., 1984; Curie et al., 2001; Jeong and Guerinot, 2009; Nozoye et al., 2015). The high stability of Fe³⁺-PS complexes allows grasses to mobilize Fe in alkaline and calcareous soils where reduction-based uptake is inefficient (Marschner et al., 1986; Römheld and Marschner, 1986; Guerinot and Yi, 1994). Some plants, such as rice, combine elements of both strategies, illustrating that plant Fe acquisition is physiologically flexible rather than strictly dichotomous (Ishimaru et al., 2006; Wairich et al., 2019; Chao and Chao, 2022).

YSL proteins belong to the Oligopeptide Transporter (OPT) superfamily together with Peptide Transporters (PTs) (Kobayashi and Nishizawa, 2012), but the evolutionary relationship between these subfamilies remains unresolved (Stacey et al., 2008; Pike et al., 2009; Cao et al., 2011; Gomolplitinant and Saier, 2011; Lubkowitz, 2011; Liu et al., 2012). Several characterized YSLs transport metal-nicotianamine or metal-phytosiderophore complexes, including Fe³⁺-PS uptake transporters in grasses. However, many YSL clades remain functionally uncharacterized, so metal or chelate transport cannot be assumed for all YSL proteins. PTs have distinct substrate ranges and evolutionary affinities; although some participate in metal homeostasis, they are not components of the canonical Strategy II uptake pathway. NAS enzymes synthesize NA, the central chelator for long-distance metal movement in plants and the precursor of phytosiderophores in grasses. Recent evidence indicates that NAS genes entered plant genomes through horizontal gene transfer (HGT) (Dirick et al., 2025), raising a key evolutionary question: did YSL transport and NAS-dependent chelation arise together, or were they assembled at different times from independent sources?

Here, we asked when YSLs became established in green plants, whether YSLs and PTs share a common evolutionary origin, when NAS-dependent chelation was added, and how YSL expansion in grasses relates to canonical Strategy II. To address these questions, we analyzed the distribution, phylogeny, and predicted structures of YSL, PT, and NAS proteins across Archaeplastida and a broad set of non-plant homologs. Our results support a modular model in which YSLs were established early in green plant evolution, PTs entered plants through an independent fungal-associated HGT route, and NAS was added later through separate fungal-to-plant transfers into mosses and euphyllophytes. This temporal uncoupling excludes plant-produced nicotianamine as a universal ancestral partner of YSL transport and indicates that the complete Strategy II pathway could emerge only after YSL transport was integrated with NAS-dependent chelator synthesis, downstream phytosiderophore biosynthesis, and phytosiderophore efflux in grasses.

## Materials and Methods

### Datasets

Complete proteomes from 66 representative species spanning major Archaeplastida lineages were compiled from JGI Phytozome (v13) (Goodstein et al., 2012) and NCBI Datasets (National Center for Biotechnology Information; https://www.ncbi.nlm.nih.gov/datasets/) (O’Leary et al., 2024) (Table S1). To investigate deeper evolutionary origins, non-plant protein sequences, including fungal, bacterial, archaeal, and other eukaryotic homologs, were downloaded locally as FASTA files from the NCBI non-redundant (NR) protein database (February 2024; https://ftp.ncbi.nlm.nih.gov/blast/db/FASTA/).

### Identification of plant OPT and NAS genes

Putative OPT and NAS homologs were identified in the Archaeplastida dataset using HMMSEARCH (E-value < e-10) from HMMER (v3.3.2) (Potter et al., 2018). Hidden Markov Model profiles for OPT (PF03169.18) and NAS (PF03059.21) were retrieved from Pfam (v35.0) (Paysan-Lafosse et al., 2025). Retrieved sequences were scanned with InterProScan (v5.69) (Jones et al., 2014) to confirm domain architecture, inspect domain position within multiple alignments, and exclude sequences containing unrelated additional domains.

Candidate proteins were filtered by length, retaining only sequences ranging from 70% to 150% of the shortest and longest *Arabidopsis* OPT or NAS proteins, respectively, to remove likely pseudogenes, protein fragments, and atypically large proteins lacking complete domain architectures. For NAS, sequences containing internal tandem domain duplications were initially removed. Because such NAS-like genes have been reported in maize and can be transcriptionally expressed despite catalytic inactivity (Zhou et al., 2013; Bonneau et al., 2016), they were reintroduced into the final dataset when detected in maize and other species (Sobic.001G395900.1.p from *Sorghum bicolor*; zea_may00001d028887_P001, zea_may00001d028888_P001, and zea_may00001d047639_P002 from *Zea mays*; and Cypan_028673 and Cypan_028675 from *Cycas panzhihuaensis*). For each gene, only the longest protein isoform was retained, and identical sequences were removed using SKIPREDUNDANT from EMBOSS (v6.6.0.0) (Rice et al., 2000).

### Statistical analyses

Statistical analyses and data visualization were performed in R (v3.6.2) and RStudio (v2025.05.1) (RStudio Team, 2020; R Core Team, 2021). Raw gene counts for OPT, PT, YSL, and NAS families, together with their genomic frequencies (number of detected homologs divided by the total number of predicted protein-coding genes per genome; Table S2), were compared among taxonomic groups using nonparametric Wilcoxon tests. To account for phylogenetic non-independence, a species tree was reconstructed from complete proteomes using OrthoFinder (v2.5.5) (Emms and Kelly, 2019), and Phylogenetic Generalized Least Squares (PGLS) models were fitted using the nlme package (v3.1-168) (Pinheiro et al., 1999). Models used log(x+1)-transformed gene counts or genomic frequencies as response variables, taxonomic group as predictor, and a Brownian-motion correlation structure (corBrownian()) derived from the species tree. Model significance was assessed by ANOVA, and post hoc pairwise contrasts were estimated as marginal means with Tukey correction using emmeans (v1.11.2-8) (Searle et al., 1980). Results are summarized in Table S3.

To test whether gene-family expansions were associated with genome-wide changes, we computed Phylogenetically Independent Contrasts (PICs) from the reconstructed species tree using the *pic()* function of the ape package (v5.8-1) (Paradis and Schliep, 2019). For each group, correlations among gene counts, genome size, and protein-coding gene number were evaluated using Pearson correlation and linear regression forced through the origin (y ∼ x + 0), ensuring statistical independence among contrasts. Regression coefficients, R² values, and P values were visualized as scatterplots (Fig. 1; Fig. S3) and are reported in Table S4. Additional R package information is provided in Methods S1.

**Figure 1.**
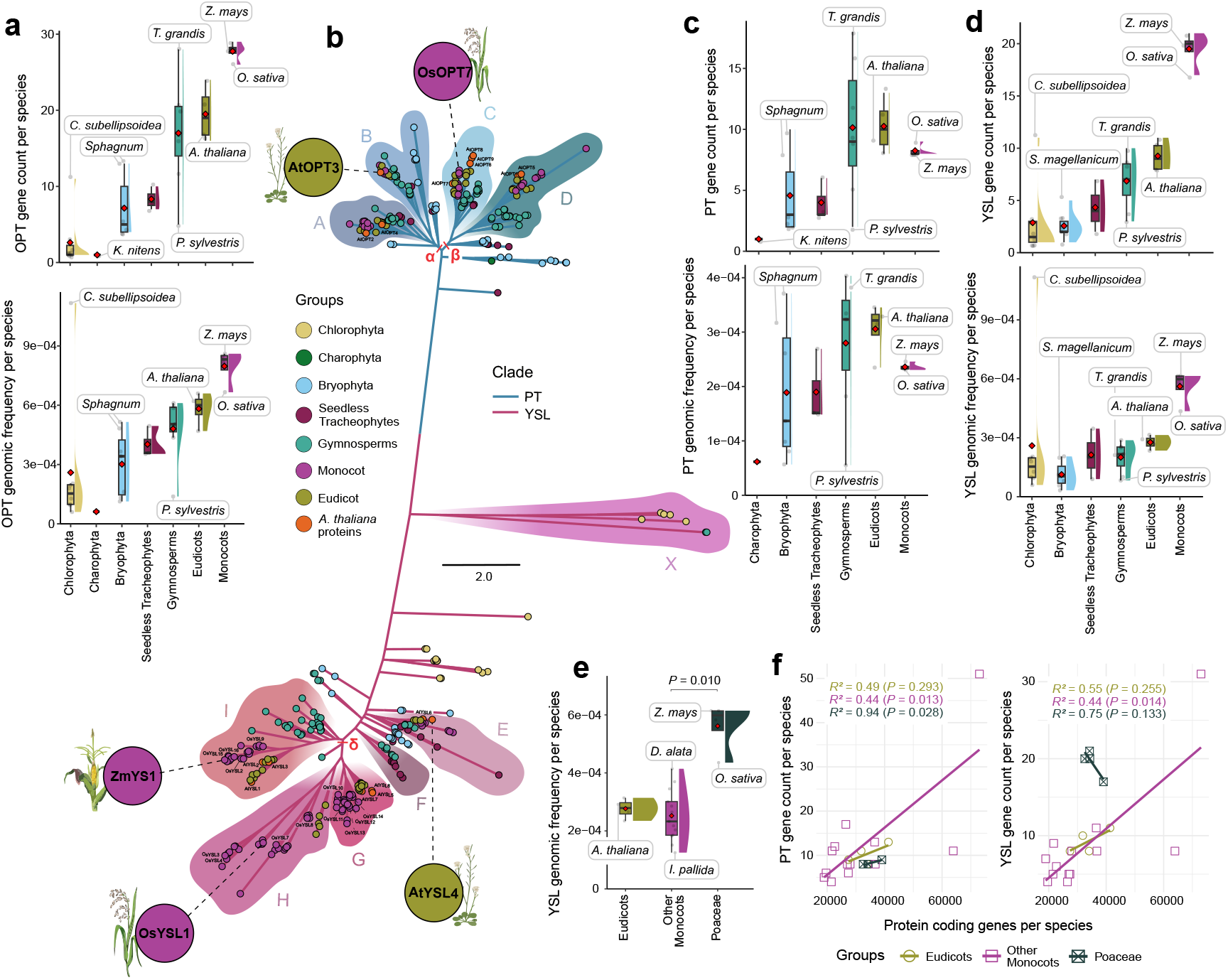
Poaceae-specific retention of YSL duplicates expanded the transporter repertoire in the lineage where Strategy II evolved. a) Gene counts (top) and normalized genomic frequencies (bottom; gene count divided by total predicted protein-coding genes) for the OPT superfamily across representative Archaeplastida genomes. b) Maximum-likelihood phylogeny of plant OPT proteins, resolving PT and YSL as deeply divergent subfamilies. Reference proteins from *Arabidopsis thaliana*, maize (*Zea mays*), and rice (*Oryza sativa*) are highlighted. Clades A– I denote orthogroups defined in this study; Greek letters indicate major ancestral expansion nodes. The complete tree with branch support values and protein identifiers is provided in Supplementary Fig. 1. c-d) Gene counts and normalized genomic frequencies for PT c) and YSL d) proteins, showing that the increase in OPT abundance across land plants is driven primarily by YSLs rather than by PTs. e) Normalized YSL genomic frequency across angiosperm groups, revealing significant enrichment in Poaceae compared with other monocots. f) Relationships between PT or YSL gene counts and total protein-coding gene number across eudicots, non-Poaceae monocots, and Poaceae. Phylogenetically informed analyses indicate that YSL accumulation in Poaceae cannot be explained by genome-wide gene content alone. In boxplots, central lines indicate medians, boxes indicate interquartile ranges, whiskers indicate 1.5× IQR, red diamonds indicate means, and half-violins show data distributions. The horizontal bar denotes a statistically significant contrast; full Wilcoxon and PGLS results are provided in Table S3, and PIC analyses are summarized in Table S4.

### Sequence alignments and phylogenetic analyses

Protein sequences were aligned with MAFFT (v7.471) (Katoh, 2002) using the parameters *--reorder --leavegappyregion --maxiterate 1000 --retree 1 --localpair --anysymbol*. Maximum-likelihood phylogenetic analyses were performed with IQ-TREE (v2.0.7) (Nguyen et al., 2015). Best-fit amino acid substitution models were selected with ModelFinder (Kalyaanamoorthy et al., 2017) using the Akaike Information Criterion (AIC) or Bayesian Information Criterion (BIC), as appropriate; selected models are listed in Table S7. Initial trees were generated with the BIONJ algorithm and optimized by Nearest Neighbor Interchange (NNI) topology searches. Branch support was estimated using the Shimodaira-Hasegawa approximate likelihood-ratio test (SH-aLRT) with 1,000 pseudoreplicates (Guindon et al., 2010). Trees were midpoint-rooted, except for PT- and YSL-specific phylogenies (Fig. 2; Figs. S5 and S6), which were rooted using representatives of the opposite subfamily as outgroups. Trees were visualized in RStudio (v2025.05.1) (RStudio Team, 2020) and FigTree (v1.4.4; http://tree.bio.ed.ac.uk/software/figtree/) and finalized in Adobe Illustrator and CorelDRAW.

**Figure 2.**
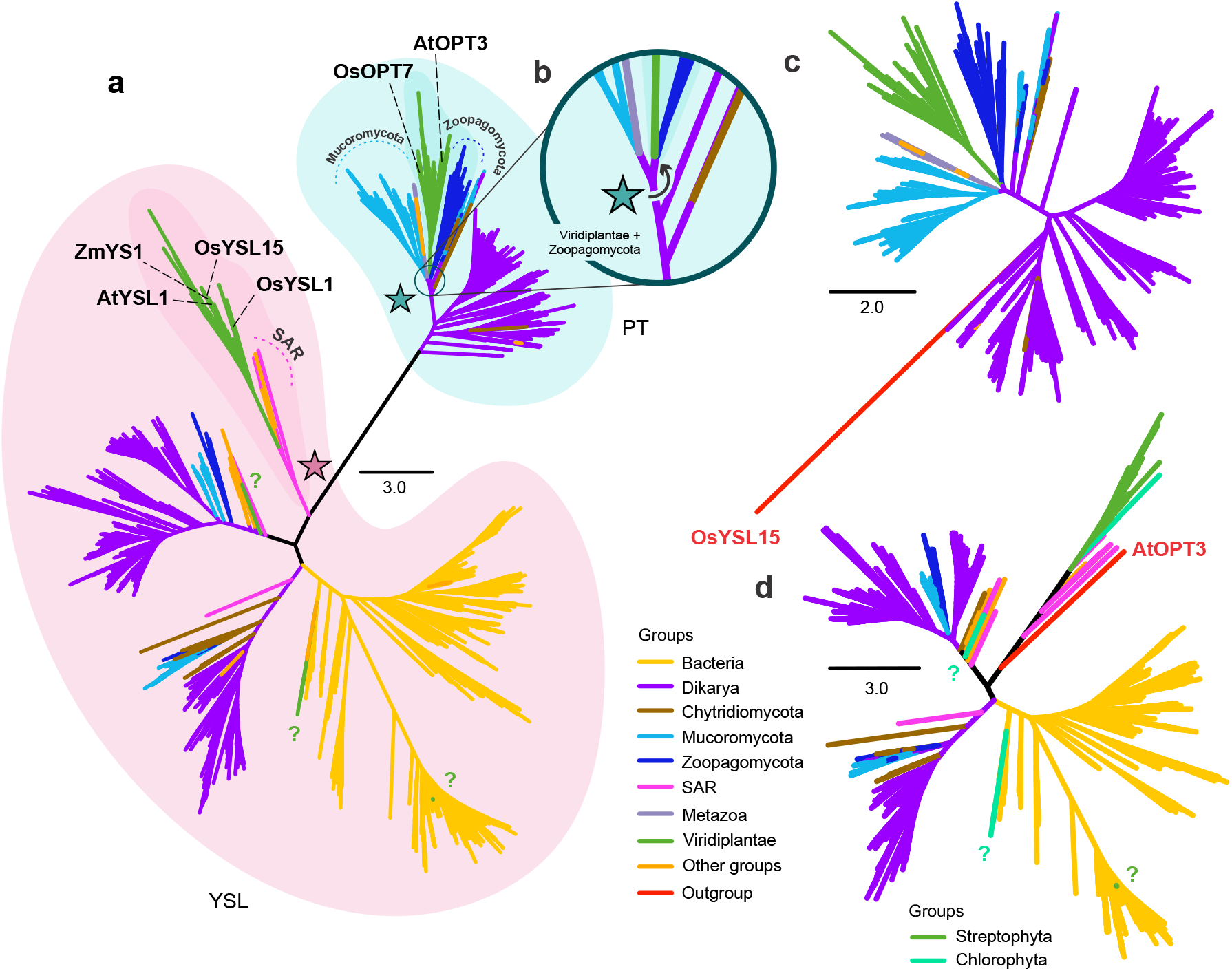
Deep phylogenies support independent evolutionary histories for PT and YSL transporters. a) Midpoint-rooted maximum-likelihood phylogeny of OPT homologs sampled from Bacteria, fungi, SAR lineages, Viridiplantae, and other eukaryotic groups. The tree resolves two major plant-associated lineages corresponding to PT and YSL proteins, with reference YSLs from Arabidopsis thaliana, maize (*Zea mays*), and rice (*Oryza sativa*) highlighted. b) Enlarged view of the plant PT placement, showing Viridiplantae PTs as sister to Zoopagomycota within a broader fungal-associated clade. c-d) Subfamily-specific phylogenies for PT c) and YSL d), each rooted with representatives of the opposite subfamily, further supporting distinct evolutionary trajectories for the two transporter lineages. Stars mark the inferred entry points of plant-associated PT and YSL lineages. Question marks indicate divergent chlorophyte YSL-like proteins with unstable or fragmented placement within the broader YSL radiation. Branch colors denote major taxonomic groups, and scale bars indicate amino acid substitutions per site. Complete cladograms with SH- aLRT support values and protein identifiers are provided in Figs. S4–S6.

### Inference of horizontal gene transfer events

Potential homologs outside plants were identified by querying the NCBI NR database with the OPT (PF03169.18) and NAS (PF03059.21) HMM profiles (E-value < 1e-30). Candidate sequences were analyzed with InterProScan (v5.69) (Jones et al., 2014) to confirm domain integrity and architecture, and the same curation criteria used for plant sequences were applied to remove partial, redundant, pseudogenic, or multidomain sequences before alignment and phylogenetic inference. For the OPT family, the initial HMM search yielded more than 90,000 putative homologs across diverse organisms. To reduce dataset size while preserving phylogenetic diversity, sequences were clustered with DIAMOND DeepClust (Buchfink et al., 2015; Buchfink et al., 2023) at 90% identity, yielding 29,529 sequences. A subsequent BLASTp search (E-value < 1e-10) (Altschul et al., 1990) was performed using all 405 Archaeplastida OPT proteins as queries, recovering 5,059 unique non-plant homologs, which were then combined with our Archaeplastida sequences for downstream phylogenetic analyses.

Candidate HGT events were inferred by comparing OPT and NAS gene trees with the species tree reconstructed from representative Archaeplastida proteomes using OrthoFinder (v2.5.5) (Emms and Kelly, 2019) (Fig. S10). Evidence for HGT included phylogenetic incongruence between gene and species trees, patchy gene distributions across lineages, and unexpectedly close sequence or structural affinity between distant taxa (Aubin et al., 2021; Mariault et al., 2025). Putative donor groups and transfer directionality were inferred from the closest non-plant clades associated with plant sequences, while considering taxonomic relationships, divergence-time constraints, and the known limitations of deep phylogenies.

Because NAS showed a patchy distribution in early-diverging plant genomes, we extended the analysis to transcriptomic datasets. Assemblies from bryophytes (hornworts, liverworts, and mosses) and lycophytes (Isoetales, Lycopodiales, and Selaginellales) were downloaded from the 1,000 Plants (1KP) project (One Thousand Plant Transcriptomes Initiative, 2019) (Table S5). BLASTp searches (E-value < e-5) were performed using all *A. thaliana* and *O. sativa* NAS sequences as queries. *A. thaliana* Actin-2 (AT3G18780) was used as a positive control for transcriptome recovery.

### Structural analysis

To compare the structural consequences of the phylogenetic divergences observed for PT, YSL, and NAS proteins, we modeled representative three-dimensional structures across selected taxa. Proteins were first analyzed with SignalP (v6.0; https://services.healthtech.dtu.dk/services/SignalP-6.0/) (Nielsen et al., 2024) to identify signal peptides or cleavage sites that could interfere with structural modeling. DeepTMHMM (v1.0.44; https://dtu.biolib.com/DeepTMHMM) (Hallgren et al., 2022) was used to predict and characterize OPT transmembrane topologies.

Protein structures were predicted with AlphaFold2 (Jumper et al., 2021) using the ColabFold Notebook pipeline (v1.5.5; https://github.com/sokrypton/colabfold) (Mirdita et al., 2022), with default settings and five relaxation iterations. Predicted structures were visualized and pairwise aligned in PyMOL (v3.1.6.1) (Schrödinger, 2015) using the align plugin. Root-mean-square deviation (RMSD) values were calculated from alpha-carbon (Cα) positions for representative OPT and NAS proteins.

Because YSL transporters undergo conformational changes associated with a proton-coupled elevator-like transport mechanism (Yamagata et al., 2022), barley (*Hordeum vulgare*) HvYS1 was remodeled and used as a structural control. All predicted OPT models were aligned against the experimental HvYS1 cryo-electron microscopy structure (PDB ID: 7WSR) (Yamagata et al., 2022). In this cryo-EM structure, the first 47 residues and the loop between transmembrane helices 6 and 7 are unresolved due to structural flexibility. Corresponding regions were manually removed from all computational models before pairwise comparisons to avoid including potentially unreliable AlphaFold-predicted flexible segments. The exact trimmed coordinates are provided in Table S8.

### NAS Sequence Similarity Network (SSN)

A NAS sequence similarity network was constructed to complement phylogenetic inference with a model-independent representation of sequence similarities. Pairwise similarities among NAS proteins were calculated with the Enzymatic Similarity Tool (EFI-EST; https://efi.igb.illinois.edu/efi-est/) (Zallot et al., 2019) using an alignment score threshold of 7, corresponding to the minimum similarity required to retain an edge. The network was visualized and refined in Cytoscape (v3.8.2) (Shannon, 2003) using the Organic Layout algorithm (Wiese et al., 2002), empirically retaining only edges connecting proteins with >51.5% sequence identity.

## Results

### Poaceae preferentially retained an expanded YSL repertoire

The OPT superfamily comprises two deeply divergent protein subfamilies, PT and YSL (Lubkowitz, 2011). Because the acronym “OPT” has historically been used both for the entire superfamily and for PT-like proteins, we use OPT here only for the superfamily and refer to the two subfamilies as PT and YSL, while retaining established gene names such as AtOPT3 when discussing individual PT genes (Stacey et al., 2008). We searched for OPT homologs in high-quality complete genomes representing major Archaeplastida lineages (Table S1) and identified 405 homologs across 34 species (Table S2). No OPT genes were detected in Rhodophyta or in the glaucophyte genome, indicating that this superfamily was either absent from the last common ancestor of Archaeplastida or was subsequently lost before becoming established in Viridiplantae (Fig. 1a). Within Viridiplantae, OPT homologs occurred at low copy number in several chlorophytes and were consistently present in all analyzed land plants.

In land plants, OPT copy number followed a clear phylogenetic pattern. Bryophytes showed the lowest median value (five genes per species), seedless tracheophytes and gymnosperms showed intermediate medians (8 and 17 genes per species, respectively), and angiosperms showed higher values, with eudicots reaching a median of 19 genes and monocots reaching the highest median of 28 genes per species. Among the six surveyed charophytes, the streptophyte algal grade most closely related to land plants, only the terrestrial *Klebsormidium* genome encoded a detectable OPT homolog (Fig. 1a). This patchy distribution is compatible with early acquisition followed by lineage-specific losses, but it also indicates that the establishment of the OPT superfamily was uneven across early streptophyte evolution.

Because genome size, protein-coding gene number, and whole-genome duplication history differ markedly among these lineages, we normalized OPT counts by the total number of predicted protein-coding genes in each genome. This genomic-frequency analysis showed a progressive increase in the fraction of gene space occupied by OPT genes from early branching chlorophytes to angiosperms, with monocots again showing the highest values (Fig. 1a). Thus, the observed pattern is not explained solely by larger genomes or increased total gene counts but also reflects lineage-specific retention or expansion of OPT homologs.

Maximum-likelihood phylogenetic analysis resolved two major clades corresponding to PT and YSL proteins (Fig. 1b; Fig. S1), consistent with previous reports (Yen et al., 2001; Gomolplitinant and Saier, 2011; Lubkowitz, 2011; Wairich et al., 2019). Each subfamily was further organized into clusters of putative orthologs, designated A-D for PT and E-I for YSL. The YSL clade also contained a set of highly divergent chlorophyte sequences that branched apart from the main plant YSL radiation and were assigned to a distinct X subclade (Fig. 1b). The distribution of PT and YSL across early-diverging Viridiplantae was irregular. One single charophyte homolog clustered with PTs and chlorophyte homologs were restricted to YSL-related clades, while land plants retained both subfamilies. Together with the absence of OPTs from several sampled algae, including *Chara braunii* and *Chlamydomonas reinhardtii*, this pattern departs from a simple model of uninterrupted vertical inheritance and suggests that the origins of PT and YSL were temporally decoupled during green plant evolution.

We then recalculated gene counts and normalized genomic frequencies separately for PT and YSL proteins (Figs. 1c-d). This separation revealed that the apparent increase in OPT abundance across land plants is driven mainly by YSLs. PT genomic frequency remained comparatively stable across land plants (Fig. 1c), whereas YSL frequency increased sharply, especially in monocots (Fig. 1d). Monocots showed a lower PT genomic frequency than eudicots but an approximately two-fold higher YSL genomic frequency. These contrasting trajectories indicate that PT and YSL repertoires evolved under different constraints. Thus, YSL expansion, rather than generalized OPT amplification, is the genomic signal associated with the grass lineage in which canonical Strategy II emerged.

Because rice and maize use YSL proteins for Fe³⁺-DMA uptake (Inoue et al., 2009), we expanded sampling to 14 additional monocot proteomes to determine whether YSL enrichment is a general monocot feature or a grass-specific pattern (Table S2). A monocot-specific OPT phylogeny recovered highly duplicated PT and YSL repertoires that remained clearly segregated (Fig. S2). Across monocots, total OPT copy number ranged from nine genes in the aquatic species *Zostera marina* to more than 80 genes in the hexaploid genome of ginger (*Zingiber officinale*). However, normalized genomic frequencies showed that YSL expansion is concentrated in Poaceae rather than shared by monocots broadly (Fig. 1e). This Poaceae-specific enrichment was the only statistically significant family-level signal, whereas PT frequencies did not differ significantly among monocot groups (Fig. 1e; Fig. S3a-b; Table S3).

Phylogenetically Independent Contrasts (Felsenstein, 1985) further showed that YSL accumulation in Poaceae is decoupled from overall genome expansion and total protein-coding gene number (Fig. 1f; Fig. S3c-d; Table S4). Whole-genome duplication alone therefore cannot account for the disproportionate enrichment of YSLs in grasses. Several YSL orthogroups, including I, H, and especially G (Fig. 1b), are markedly expanded in Poaceae, consistent with preferential retention of duplicates arising from both whole-genome and small-scale gene duplication events. These results identify a grass-specific expansion of the YSL repertoire associated with Strategy II evolution, but they do not establish that all expanded orthogroups function as Fe-phytosiderophore transporters.

### YSL and PT proteins entered the green plant gene pool through independent evolutionary routes

The discontinuous distribution of OPT homologs across Archaeplastida raised the possibility that PT and YSL proteins have distinct origins. To test this, we searched the NCBI non-redundant database and recovered 91,353 candidate non-plant OPT homologs. After clustering and filtering to reduce redundancy, while preserving phylogenetic breadth, we reconstructed a deep maximum-likelihood phylogeny including 5,059 representative homologs from bacteria, archaea, diverse eukaryotes, and our Archaeplastida dataset. The tree resolved major fungal groups, a bacterial clade, two major plant-associated clades, and additional eukaryotic lineages, including Haptista, Discoba, Amoebozoa, Archaea, and the SAR supergroup (Stramenopila, Alveolata, and Rhizaria) (Richards et al., 2009) (Fig. 2a; Fig. S4). Within this broader tree, plant PT and YSL formed deeply separated lineages. Streptophytes PTs were most closely affiliated with fungal homologs (Fig. 2b), whereas Viridiplantae YSLs were associated with a distinct assemblage including Haptista and SAR members such as oomycetes and ochrophytes (Fig. 2a; SH-aLRT support values >79%). These phylogenetic relationships indicate that PT and YSL do not represent a single plant origin followed by subfamily divergence and functional diversification.

To refine these inferences, we reconstructed PT- and YSL-specific phylogenies, rooting each subfamily with representatives of the other. In the PT tree, homologs from *Klebsormidium nitens* and land plants formed a well-supported sister relationship with Zoopagomycota within a broader Mucoromycota-associated clade (SH-aLRT support values >94%; Fig. 2c; Fig. S5). This topology supports a fungal-associated origin for plant PT repertoires. Because early-diverging Mucoromycota and related fungi are central to ancient plant-fungal associations (Richards et al., 2009; Bidartondo et al., 2011; Feijen et al., 2018), the PT pattern is consistent with acquisition during the early history of streptophytes, probably in the context of intimate plant-fungal contact. Importantly, this origin is independent from the evolutionary history inferred for YSLs.

By contrast, Viridiplantae YSLs showed an older and more complex evolutionary pattern (Fig. 2d; Fig. S6). Most Viridiplantae YSLs formed a monophyletic clade associated with SAR lineages (SH-aLRT = 91.1%), whereas the divergent chlorophyte X proteins were fragmented across the tree. Mamiellophyceae homologs formed a strongly supported clade with Discoba (SH-aLRT = 99.6%) and branched near bacterial sequences, while homologs from *Coccomyxa subellipsoidea* and *Chromochloris zofingiensis* occurred either near Viridiplantae or within a broader eukaryotic assemblage including Amoebozoa, Haptista, and Oomycota (SH-aLRT support values >80%). These placements are consistent with repeated ancient transfers and losses, but tree topology alone cannot establish the direction of the earliest events. The deep separation of YSL from PT, its proximity to bacterial-associated lineages, and its broad eukaryotic distribution are compatible with a prokaryotic ancestry and early establishment in green plants. However, the data do not distinguish whether the principal Viridiplantae YSL lineage originated within early green plants or was acquired from another ancient lineage. In either case, plant YSLs clearly predate streptophyte terrestrialization and NAS-dependent phytosiderophore biosynthesis.

### YSL proteins form a structurally conserved lineage distinct from PT transporters

The phylogenetic separation between PT and YSL proteins predicts structural differences, despite belonging to the same superfamily and sharing transmembrane architecture. We therefore compared high-confidence structural models across representative taxa (Fig. 3; Fig. S7), using both AlphaFold2-predicted and cryo-electron microscopy structures of barley (*Hordeum vulgare*) YELLOW STRIPE 1 (HvYS1) (Yamagata et al., 2022) as controls. The AlphaFold2 HvYS1 model agreed closely with the experimental cryo-EM structure, supporting the use of these models for comparative analyses. All-versus-all structural alignments revealed strong conservation within each subfamily (RMSD <2.5 Å for PT-PT and YSL-YSL comparisons), but pronounced divergence between PT and YSL proteins, with average RMSD values exceeding 8.5 Å (Fig. 3a). Although individual exceptions occurred, such as the oomycete YSL from *Achlya hypogyna* showing partial structural similarity to land plant PTs, the overall structural dendrogram clearly separated PT and YSL proteins.

**Figure 3.**
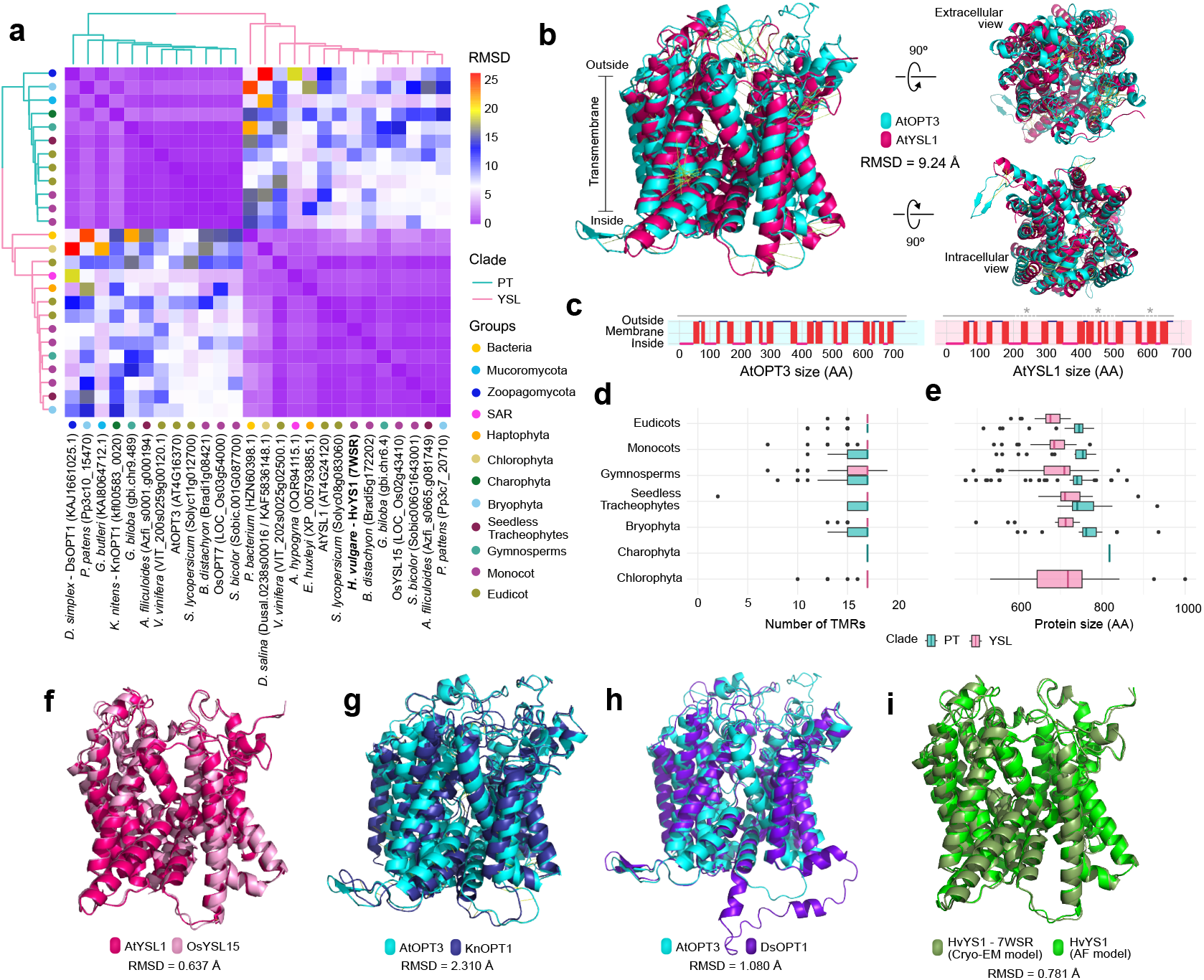
Structural comparisons distinguish YSL proteins from PT transporters. a) Pairwise structural similarity among representative OPT protein models, shown as a heatmap of RMSD values in Å. Hierarchical clustering separates most PT and YSL proteins, indicating strong structural conservation within each subfamily and pronounced divergence between subfamilies. b) Structural alignment of *Arabidopsis thaliana* OPT3 and YSL1 from lateral and intracellular/extracellular views, illustrating differences in helix packing, loop architecture, and channel organization despite a shared transmembrane core. c) Predicted transmembrane topologies of AtOPT3 and AtYSL1 generated with DeepTMHMM; blue, magenta, and red indicate extracellular loops, intracellular regions, and transmembrane helices, respectively. d-e) Number of predicted transmembrane regions d) and total protein length e) across representative plant groups for PT and YSL proteins. Central lines indicate medians, boxes indicate interquartile ranges, whiskers indicate 1.5× IQR, and black dots indicate outliers. f–i) Representative pairwise structural alignments showing conservation within YSLs (AtYSL1 and rice OsYSL15; f), structural similarity between AtOPT3 and PT homologs from *Klebsormidium nitens* (KnOPT1; g) and the fungus Dispira simplex (DsOPT1; h), and agreement between the predicted barley HvYS1 model and the experimental HvYS1 cryo-EM structure (PDB ID: 7WSR; i). RMSD values are shown for each comparison.

Direct comparison between *Arabidopsis thaliana* OPT3 and YSL1 illustrates the structural basis of this separation (Fig. 3b). The transmembrane backbone is broadly conserved, but PT and YSL proteins differ in helix arrangement, loop architecture, and the organization of the internal channel. Transmembrane topology predictions further indicate distinct membrane-crossing patterns (Fig. 3c). AtOPT3 contains longer extracellular loops and a more continuous alternating arrangement of transmembrane helices, whereas AtYSL1 shows localized disruptions in this arrangement and shorter extracellular loops. PT proteins also tend to have longer and more variable N- and C-terminal regions, whereas YSL proteins are more compact and conserved, especially near the C-terminus. These differences are concentrated in regions likely to influence substrate access, gating, and conformational transitions rather than in the existence of the transmembrane core itself.

Across Archaeplastida, PT and YSL proteins showed similar numbers of predicted transmembrane regions (Fig. 3d), but PTs were consistently longer (Fig. 3e). This agrees with previous reports that OPT subfamilies share only 10-16% sequence similarity (Yen et al., 2001; Stacey et al., 2008; Lubkowitz, 2011) and reinforces the conclusion that plant PT proteins are evolutionarily closer to fungal homologs than to plant YSLs. YSL proteins maintained a highly conserved global architecture across distant taxa (Fig. 3f; Fig. S7a), whereas the *Arabidopsis* AtOPT3 was structurally closer to PT homologs from divergent lineages, including algae and fungi, than to plant YSL1 (Fig. 3g-h; Fig. S7b-c). The high agreement between predicted and experimental HvYS1 structures provides an internal control for these comparisons (Fig. 3i; Fig. S7d).

Together, the structural and phylogenetic evidence indicates that PT and YSL proteins are not interchangeable branches of a recently diversified plant family. Rather, they are structurally distinct lineages that entered green plant evolution through different routes and subsequently evolved under different constraints. For the Strategy II question, YSL provides the relevant pre-existing transporter repertoire, whereas PTs show that the plant OPT superfamily is a composite evolutionary assemblage. Direct functional analyses are still required to determine the substrates of ancestral YSLs and of the YSL proteins belonging to the orthogroups preferentially expanded in grasses.

### Independent fungal-to-plant HGT events introduced NAS into mosses and euphyllophytes

NAS enzymes synthesize NA, which functions as a metal chelator in plants and as the precursor for phytosiderophore biosynthesis in Strategy II (Fig. 4a). If YSL-mediated transport were ancestrally coupled to NA, NAS genes would be expected to broadly co-occur with YSL genes across early plant lineages. We therefore examined the distribution and evolutionary history of NAS to determine whether the chelator-synthesis module appeared together with, or after, the YSL transport module.

**Figure 4.**
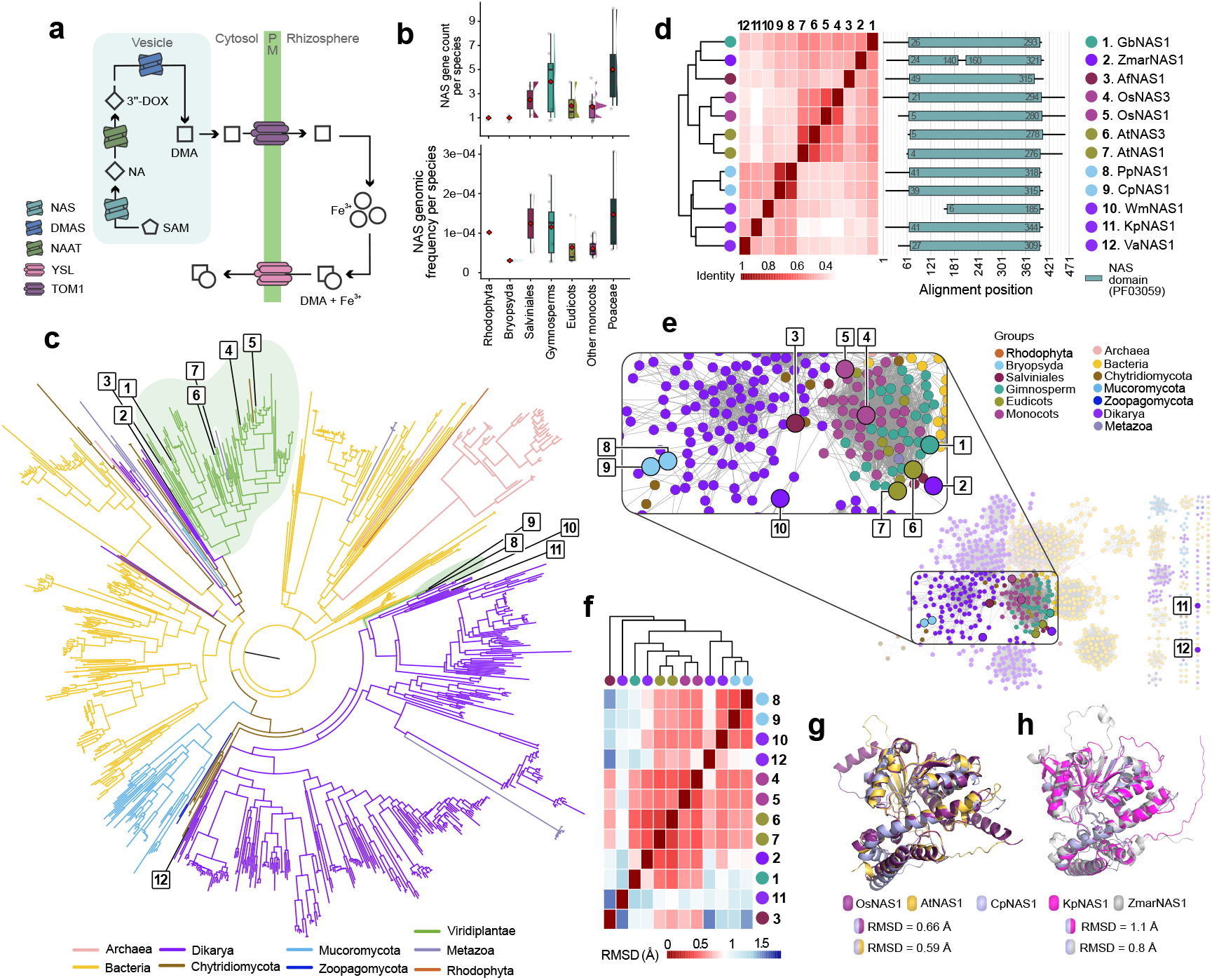
Independent fungal-to-plant HGT events introduced NAS into mosses and euphyllophytes. a) Schematic representation of Strategy II iron uptake, highlighting the role of NAS in phytosiderophore biosynthesis. NAS converts S-adenosylmethionine (SAM) into nicotianamine (NA), which is subsequently converted by NAAT and DMAS into deoxymugineic acid (DMA). TOM1 exports DMA to the rhizosphere, where it chelates Fe³⁺, and YSL transporters import Fe³⁺-DMA complexes across the plasma membrane (PM). b) NAS gene counts (top) and normalized genomic frequencies (bottom; gene count divided by total predicted protein-coding genes) across major Archaeplastida lineages. Central lines indicate medians, boxes indicate interquartile ranges, whiskers indicate 1.5× IQR, red diamonds indicate means, and half-violins show data distributions. c) Midpoint-rooted maximum-likelihood phylogeny of NAS proteins sampled across bacteria, fungi, plants, animals, and other lineages. Plant NAS proteins do not form a single monophyletic group; instead, moss and euphyllophyte sequences cluster with distinct fungal lineages, supporting independent fungal-to-plant HGT events. d) Pairwise sequence identity among selected NAS proteins from bryophytes, euphyllophytes, and fungi. The domain schematic on the right shows the position of the NAS domain identified by InterProScan (Table S6). e) Sequence similarity network of NAS proteins, showing separation between euphyllophyte NAS proteins and bryophyte/fungal-associated sequences. Node colors denote taxonomic groups, and numbered nodes correspond to the proteins highlighted in (d). f) Pairwise structural similarity among selected NAS models, shown as a heatmap of RMSD values in Å. g-h) Representative structural alignments comparing *Ceratodon purpureus* NAS1 (CpNAS1) with angiosperm NAS proteins (AtNAS1 and OsNAS1; g) and with the fungal *Zalerion maritima* NAS1 (ZmarNAS1; h), illustrating conservation of the NAS fold despite independent acquisitions.

In contrast to YSL, NAS showed a restricted and patchy distribution. We detected only 91 NAS homologs across the analyzed plant lineages (Table S2), distributed among Rhodophyta (1), Bryopsida (2), Salviniales (5), gymnosperms (30), and angiosperms (53) (Fig. 4b). In Poaceae, NAS genes were organized into two functionally distinct paralogous subclades (Fig. S8): Clade 1, including rice OsNAS1/2, associated with root Fe uptake; and Clade 2, including OsNAS3, associated with shoot Fe homeostasis (Inoue et al., 2003; Mizuno et al., 2003; Bonneau et al., 2016; Seregin and Kozhevnikova, 2023). Despite these lineage-specific duplications, NAS gene counts and normalized genomic frequencies remained low overall and did not differ significantly among major Archaeplastida groups (Fig. 4b; Table S3). The absence of NAS from several early-diverging lineages that nevertheless possess YSLs led us to examine transcriptomic evidence from bryophytes and lycophytes. Searches of 1KP transcriptomes (One Thousand Plant Transcriptomes Initiative, 2019) also indicate that NAS transcripts are absent from liverworts, hornworts, and lycophytes and are restricted to Bryopsida mosses and Euphyllophyta (Table S5).

Deep phylogenetic analysis of 791 non-plant NAS homologs further supported a complex origin. The NAS family has bacterial roots and a scattered distribution across the tree of life, with representatives in Archaea, Metazoa, fungi, and plants, consistent with previous studies (Laffont and Arnoux, 2020; Gilbert and Maumus, 2022; Dirick et al., 2025) (Fig. 4c; Fig. S9). Archaeplastida NAS proteins did not form a monophyletic clade. The single Rhodophyta NAS from *Chondrus crispus* branched independently with bacterial Thermoactinomycetaceae (SH-aLRT = 84%). Tracheophyte NAS proteins formed a sister relationship with a fungal clade dominated by estuarine taxa, including Lulworthiaceae as *Zalerion maritima*, whereas moss NAS proteins grouped with terrestrial Pezizomycetes fungi such as *Kalaharituber pfeilii* (SH-aLRT support values >94%). This topology indicates that plant NAS was not inherited from a common land plant ancestor, but was acquired independently in specific terrestrial plant lineages through fungal-to-plant HGT.

Because NAS amino acid sequences appear superficially conserved, we tested whether phylogenetic separation was also recovered by model-independent comparisons. Pairwise identity analyses showed that the core NAS domain is conserved, but moss NAS proteins share higher primary-sequence identity with Dikarya fungi than with tracheophyte NAS proteins (Fig. 4d; Table S6). Tracheophyte NAS proteins, in turn, possess a distinct extended C-terminal region. A sequence similarity network (SSN) recovered the same separation: moss NAS proteins clustered within a Dikarya-associated network, whereas tracheophyte proteins formed a distinct plant-associated cluster (Fig. 4e). These sequence-level affinities support at least two independent plant NAS acquisitions from ecologically distinct Dikarya fungal donors, in agreement with recent evidence for repeated NAS transfers and replacements in plants (Dirick et al., 2025).

Structural modeling showed that NAS enzymes retained a strongly conserved overall fold despite their independent evolutionary histories. Pairwise structural alignments produced low RMSD values (<1.5 Å) across all selected models (Fig. 4g-h). However, the moss *Ceratodon purpureus* NAS (CpNAS1), despite its phylogenetic proximity to fungal homologs, was structurally more similar to angiosperm NAS proteins (AtNAS1 and OsNAS1; RMSD = 0.59-0.66 Å) than to fungal NAS proteins (RMSD = 0.8-1.1 Å) (Fig. 4g-h). This pattern is consistent with similar functional constraints acting on independently acquired plant NAS enzymes, rather than with inheritance from a single plant ancestor. NAS therefore represents a later chelator-synthesis module incorporated into plant metal homeostasis after YSL lineages were already established and diversified in green plants.

## Discussion

The evolution of plant Fe acquisition has usually been framed as a contrast between reduction-based Strategy I and chelation-based Strategy II (Römheld and Marschner, 1986; Kobayashi and Nishizawa, 2012; Connorton et al., 2017; Grillet and Schmidt, 2019; Wairich et al., 2019; Chao and Chao, 2022). However, the broad distribution of genes associated with metal transport and chelation complicates this dichotomy and raises three questions: which components are ancient, which are grass-specific, and how was a complete phytosiderophore-based pathway assembled? By reconstructing the histories of YSL, PT, and NAS proteins across Archaeplastida and beyond, we show that the Strategy II toolkit was not inherited as a single ancestral plant pathway. Instead, it was assembled asynchronously: YSLs constitute the oldest transporter lineage later recruited into Strategy II, NAS was added through independent fungal-to-plant HGT events, and PTs reveal that the plant OPT superfamily itself is an evolutionary mosaic of distinct protein lineages (Fig. 5).

**Figure 5.**
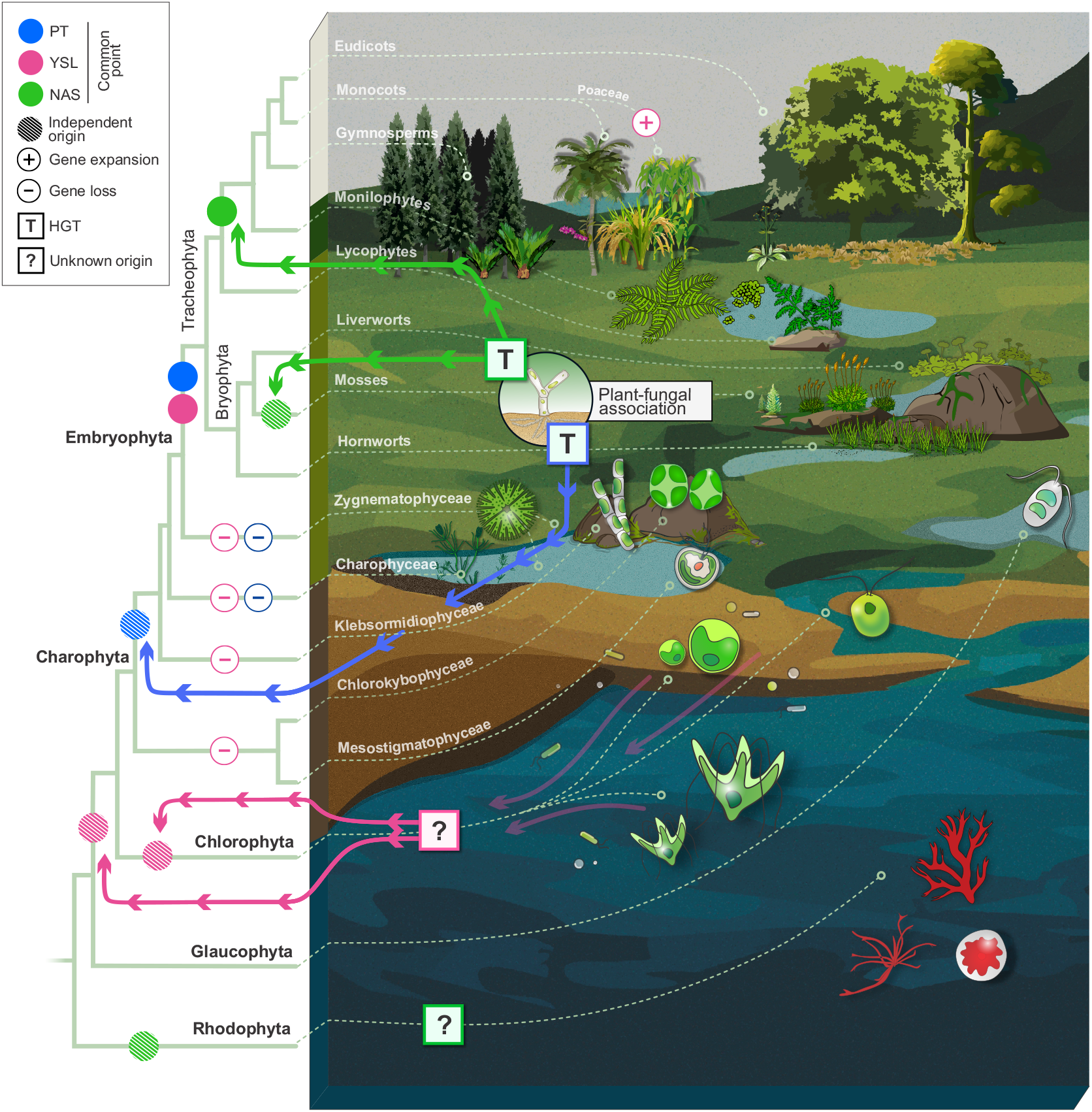
Evolutionary model for the asynchronous assembly of the Strategy II gene toolkit. The schematic integrates Archaeplastida phylogeny, the transition from aquatic to terrestrial habitats, and recurrent interactions with microbial communities. Symbols on the phylogeny indicate inferred evolutionary events for PT (blue), YSL (pink), and NAS (green): solid circles mark inferred establishment points, hatched circles indicate independent origins or unresolved placements, T-labeled squares denote horizontal gene transfer (HGT), question marks indicate uncertain origins or directionality, plus signs indicate lineage-specific gene expansion, and minus signs indicate gene loss. The model places YSL as an ancient transporter lineage established in green plants before the appearance of plant NAS, while leaving the exact origin and direction of the earliest YSL event unresolved. PT proteins entered plants independently, probably through fungal-associated HGT, and illustrate the composite origin of the plant OPT superfamily. NAS entered distinct land plant lineages later through multiple fungal-to-plant HGT events. Canonical Strategy II could emerge only after NAS-dependent chelator synthesis was integrated with NAAT/DMAS-mediated phytosiderophore biosynthesis, TOM-mediated efflux, and a YSL transporter repertoire. NAAT, DMAS, and TOM were not analyzed here; their placement indicates required pathway context rather than inferred origins or timing.

The strongest lineage-specific signal in our analysis is the expansion of YSLs in grasses. Historically, YSL transporters were first associated with the Major Facilitator Superfamily and then reclassified within the fungal OPT family alongside PT proteins (Lubkowitz et al., 1997; Curie et al., 2001; Yen et al., 2001). This classification created the impression that PT and YSL might be sister subfamilies with a shared plant origin, despite their distinct substrates and functions (Gomolplitinant and Saier, 2011; Lubkowitz, 2011; Kumar et al., 2025). Our data show that PT abundance remains relatively stable across land plants, whereas YSL abundance increases asymmetrically and culminates in a Poaceae-specific enrichment. This signal persists after phylogenetic correction and is disproportionate to total gene number, indicating preferential retention of YSL duplicates in grasses. However, this expansion cannot be equated with duplication of a grass-specific Fe³⁺-phytosiderophore uptake clade. Maize ZmYS1 and rice OsYSL15 occur within broader YSL radiations that also contain non-grass homologs (Force et al., 1999; Lynch and Conery, 2000; Curie et al., 2001; Inoue et al., 2009; Zhang et al., 2018). Preferential retention therefore likely enlarged the repertoire available for regulatory, tissue-specific, and substrate specialization in the lineage where Strategy II evolved; which expanded orthogroups contribute directly to Fe-phytosiderophore uptake remains unresolved.

From a broader evolutionary perspective, YSLs are among the oldest transporter lineages later recruited into plant chelation-based Fe acquisition. Their deepest relationships point to prokaryotic ancestry and a complex history of transfers among early eukaryotic lineages. Our analyses do not identify a single extant YSL orthogroup as ancestral, nor do they distinguish whether the principal Viridiplantae lineage originated within early green plants or was acquired from another ancient lineage. Functional evidence nevertheless establishes metal-related roles for some YSL clades. AtYSL4 and AtYSL6 localize to internal cellular membranes, and mutant analyses implicate them in Mn and Fe homeostasis (Conte et al., 2013), whereas other characterized YSLs transport metal-NA or metal-PS complexes (Curie et al., 2001; Inoue et al., 2009; Conte and Walker, 2012; Zheng et al., 2012; Singh et al., 2023; Kumar et al., 2025). These examples should not be generalized to all YSLs. The fragmented placement of chlorophyte YSL-like proteins indicates additional transfers and losses in green algae, and the substrates of the earliest plant YSLs remain unknown. Thus, our results separate the origin of YSL proteins from both the origin of PTs and the later assembly of NAS-dependent phytosiderophore biosynthesis.

The PT results are important because they prevent an overly simple interpretation of OPT evolution. Plant PTs cluster with early-diverging fungal lineages, particularly within a Mucoromycota/Zoopagomycota-associated context, consistent with acquisition during periods of close plant-fungal interaction (Richards et al., 2009; Bidartondo et al., 2011; Delaux et al., 2015; Feijen et al., 2018; Lutzoni et al., 2018; Morris et al., 2018; Bowles et al., 2023). Such events fit a broader pattern in which HGT contributed to early land plant evolution (Richards et al., 2009; Lutzoni et al., 2018; Ma et al., 2022). However, PTs should not be interpreted as Strategy II components. Their role in this study is comparative: they demonstrate that the OPT superfamily combines protein lineages with independent origins, distinct structures, and different evolutionary constraints. Recognizing this distinction sharpens the main conclusion that the YSL module, not PTs, is the relevant ancestral transporter background from which Strategy II could later evolve.

NAS followed a markedly different evolutionary trajectory. Whereas YSLs were broadly retained across green plants, NAS is absent from many early-diverging lineages and entered land plants later through multiple HGT events from Dikarya fungi. This temporal uncoupling excludes plant-produced nicotianamine and nicotinanamine-derived molecules as the universal ancestral ligand associated with YSL transport. If early YSLs participated in metal movement, they may have recognized complexes formed with other ligands, potentially including microbial siderophores or related metallophores available in early terrestrial environments. Alternatively, some ancestral YSLs may have transported substrates unrelated to metal chelation. These possibilities can be tested directly by characterizing the substrate specificity of YSL proteins from liverworts and hornworts, which lack detected NAS genes and therefore the canonical capacity to synthesize NA and NA-derived phytosiderophores. NAS itself has deep prokaryotic roots and likely originated as part of ancient metal-resistance or metal-homeostasis chemistry (Laffont and Arnoux, 2020). Consistent with this ancestry, structural studies of archaeal NAS-like enzymes show that the catalytic architecture required for nicotianamine synthesis predates plants (Dreyfus et al., 2009). The recurrent transfer of NAS-like genes across bacteria, fungi, plants, and animals further supports the view that metallophore synthesis is an operational module that can be mobilized across distant lineages (Jain et al., 1999; Soucy et al., 2015).

Fungal NAS enzymes provide a plausible immediate source for plant NAS acquisitions. In filamentous fungi such as *Neurospora crassa*, NAS genes encode functional enzymes that synthesize NA as part of intracellular metal chelation and are induced under zinc deprivation (Trampczynska et al., 2006). Thus, this biochemical capacity transferred to plants was most likely already functional in fungi. Our analyses indicate that mosses and euphyllophytes acquired NAS through different fungal donors, consistent with recent evidence for multiple independent NAS insertions and complex replacements in plant evolution (Dirick et al., 2025). The extended and autoinhibitory C-terminal region present in euphyllophyte NAS proteins provides an additional layer of functional divergence from fungal and bryophyte distant homologs (Seebach et al., 2023). This structural and phylogenetic bifurcation supports a model in which similar catalytic activities were independently integrated into different plant lineages after NAS HGT events.

The temporal gap between YSL establishment and NAS acquisition raises the central question of this study: what substrates did YSLs recognize before plant lineages acquired nicotianamine synthesis? Aquatic lineages can acquire and transport Fe through mechanisms that do not require a YSL/NAS system, including reductive uptake, permeases, and exploitation of microbial siderophores (Behnke and LaRoche, 2020; Blaby-Haas, 2022). During terrestrialization, however, plants encountered soils in which Fe was often poorly soluble, while intimate associations with fungi and bacteria became a major ecological interface (Knack et al., 2015; de Vries and Archibald, 2018; Kumar et al., 2025). Microbial metal-ligand complexes therefore represent plausible, but untested, ancestral substrates for early plant YSLs. Substrates unrelated to metal chelation remain an alternative.

The recently proposed Strategy III framework provides one plausible ecological model (Gu et al., 2025). In this model, plants exploit microbial siderophores through endocytosis, uptake by unidentified transporters, or direct transport by YS1/YSL-family proteins. Ancestral plants may therefore have used siderophores or related metallophores produced by soil microorganisms, with some YSLs mediating uptake or intracellular redistribution of the resulting metal-ligand complexes. This scenario is consistent with the deep bacterial-associated history of YSLs, the demonstrated ability of some family members to transport Fe-chelate complexes, and the predominant role of characterized YSLs in internal metal distribution in extant plants (Curie et al., 2001; Conte and Walker, 2012; Zheng et al., 2012; Zheng et al., 2012; Bowles et al., 2023; Seregin and Kozhevnikova, 2023; Singh et al., 2023; Gu et al., 2025; Kumar et al., 2025). After NAS acquisition, plants gained endogenous NA synthesis, and grasses integrated this module with phytosiderophore production and secretion. Poaceae-specific YSL expansion then broadened the repertoire from which Fe³⁺-phytosiderophore uptake, internal metal distribution, and other functions could be recruited, although the contribution of individual expanded orthogroups remains unresolved.

Together, these findings support Strategy II as a product of modular evolution rather than a single vertically inherited pathway. The pre-existing component was an ancient YSL transporter repertoire that was later coupled to plant-produced nicotianamine and phytosiderophore chemistry. NAS entered mosses and euphyllophytes through independent fungal-to-plant HGT events, while a complete Strategy II pathway additionally required NAAT, DMAS, and TOM-mediated efflux. In Poaceae, preferential retention and diversification of YSL paralogs expanded the transporter repertoire in the lineage where phytosiderophore-based Fe uptake evolved. This asynchronous assembly illustrates how a complex physiological trait can arise through the integration of components with distinct evolutionary histories and highlights plant-microbe interactions as a source of innovation during plant evolution.

## Acknowledgments

This work was supported by the Coordenação de Aperfeiçoamento de Pessoal de Nível Superior - Brazil (CAPES; Finance Code 001) and by the Fundação de Amparo à Pesquisa do Estado de Minas Gerais (FAPEMIG/Brazil), which provided PhD scholarships to M.L.C.B.C. and W.F.C.R., respectively. Computational analyses, statistics, and data processing were carried out using the Gloriosos clusters at PPG Bioinformatics (UFMG/Brazil) and the Striga server at the Laboratório de Interação Vegetal (LIVe) (ESALQ/USP). Deep phylogenetic analyses were performed using computational resources from the Del Bem Lab, Department of Genetics (LGN), ESALQ/USP, as well as LaGEEvo and CENAPAD-SP. The authors would like to thank Dr. Elsbeth Walker and Dr. Mary Lou Guerinot for reading the manuscript and providing feedback.

## Author contributions

L-ED-B conceived and designed the research. MLCBC and WFCR performed data curation, formal analysis, investigation, methodology, validation, visualization, and writing (original draft, review, and editing). MLCBC and WFCR contributed equally to this work. HFG contributed to data curation, formal analysis, investigation, methodology, and writing (original draft). L-ED-B carried out conceptualization, data curation, formal analysis, funding acquisition, project administration, resources, supervision, validation, methodology, visualization, and writing. FKR and JEL contributed to data interpretation and manuscript structure. All authors participated in manuscript editing and approved the final version.

## Data availability

All data supporting the findings of this study are available in the Supplementary Tables. Generated and analyzed datasets, including gene counts and genomic frequencies, taxonomy tables, and raw phylogenetic trees in .nwk format, have been deposited in Figshare under DOI: https://doi.org/10.6084/m9.figshare.31860574.

Custom R scripts and packages used to quantify gene distributions, perform statistical analyses, and plot phylogenies also are publicly available in the same Figshare repository.

## Supplementary Figures (short legends)

Figure S1. Maximum-likelihood phylogenetic tree of Archaeplastida OPT proteins.

Figure S2. Maximum-likelihood phylogenetic tree of monocot OPT proteins.

Figure S3. Poaceae-specific YSL expansion is disproportionate to genome-wide gene content.

Figure S4. Deep maximum-likelihood phylogenetic tree of OPT proteins.

Figure S5. Deep maximum-likelihood phylogenetic tree of the PT subfamily.

Figure S6. Deep maximum-likelihood phylogenetic tree of the YSL subfamily.

Figure S7. Structural overview of OPT proteins across species.

Figure S8. Maximum-likelihood phylogenetic tree of Archaeplastida Nicotianamine Synthases (NAS).

Figure S9. Deep maximum-likelihood phylogenetic tree of Nicotianamine Synthase (NAS). Figure S10. Consensus species phylogeny of Archaeplastida.

## Supplementary Tables (titles)

Table S1. Archaeplastida proteomes used in this study.

Table S2. Distribution and quantitative summary of OPT and NAS homologs across Archaeplastida.

Table S3. Statistical comparisons of gene-family distributions across Archaeplastida.

Table S4. Correlation and linear regression analyses based on Phylogenetically Independent Contrasts (PICs).

Table S5. Identification of Nicotianamine Synthase (NAS) homologs in bryophyte and lycophyte transcriptomes from the 1,000 Plants (1KP) project.

Table S6. NAS domain characterization and interspecies comparison.

Table S7. Evolutionary amino acid substitution models applied in IQ-TREE analyses.

Table S8. Structural trimming coordinates for three-dimensional OPT models.

