## Supporting Information for "Asynchronous origins of Yellow-Stripe transporters and Nicotianamine Synthase underlie the evolution of plant Strategy-II iron uptake"

### This PDF file includes:

List of Contents

Supporting Text

Figures S1 to S10

Tables S1 to S8

**List of Contents**

**Methods S1**

**Figure S1.** Maximum-likelihood phylogenetic tree of Archaeplastida OPT proteins.

**Figure S2.** Maximum-likelihood phylogenetic tree of monocot OPT proteins.

**Figure S3.** YSL transporters exhibit a Poaceae-specific expansion independent of genome size.

**Figure S4.** Deep maximum-likelihood phylogenetic tree of OPT proteins.

**Figure S5.** Deep maximum-likelihood phylogenetic tree of the PT subfamily.

**Figure S6.** Deep maximum-likelihood phylogenetic tree of the YSL subfamily.

**Figure S7.** Structural overview of OPT proteins across species.

**Figure S8.** Maximum-likelihood phylogenetic tree of Archaeplastida Nicotianamine Synthases (NAS).

**Figure S9.** Deep maximum-likelihood phylogenetic tree of Nicotianamine Synthase (NAS).

**Figure S10.** Consensus species phylogeny of Archaeplastida.

**Table S1.** Archaeplastida proteomes used in this study.

**Table S2.** Distribution and quantitative summary of OPT and NAS homologs across Archaeplastida.

**Table S3.** Statistical comparisons of gene-family distributions across Archaeplastida.

**Table S4.** Correlation and linear regression analyses based on Phylogenetically Independent Contrasts (PICs).

**Table S5.** Identification of Nicotianamine Synthase (NAS) homologs in bryophyte and lycophyte transcriptomes from the 1,000 Plants (1KP) project.

**Table S6.** NAS domain characterization and interspecies comparison.

**Table S7.** Evolutionary amino acid substitution models applied in IQ-TREE analyses.

**Table S8.** Structural trimming coordinates for three-dimensional OPT models.

**References**

**Methods S1**

Statistical R-packages and plotting

All data processing, statistical analyses, and figure generation were carried out in R (v3.6.2) within RStudio (v2025.05.1) (RStudio Team, 2020; R Core Team, 2021). The computational environment combined general-purpose statistical libraries with specialized packages for phylogenetic comparative analyses and graphical assembly.

Phylogenetic and statistical modeling were performed using ape (v5.8-1) (Paradis and Schliep, 2019) for tree manipulation and computation of phylogenetically independent contrasts (PICs), nlme (v3.1-168) (Pinheiro et al., 1999; Mixed-Effects Models in S and S-PLUS, 2000) for phylogenetic generalized least squares (PGLS) models, and phytools (v2.5-2) (Revell, 2024) for phylogenetic utilities such as midpoint rooting and ultrametric adjustment. Marginal means and post hoc Tukey contrasts were obtained with emmeans (v1.11.2-8) (Searle et al., 1980; Lenth et al., 2025).

Data wrangling and factor management were handled with dplyr (v1.1.4) (Wickham et al., 2025) and forcats (v1.0.1) (Wickham, 2025), while stringr (v1.5.2) (Wickham, 2023) and stringdist (v0.9.15) (Loo, 2014) supported string normalization and fuzzy matching between dataset and tree labels. The rstatix package (v0.7.2) (Kassambara, 2023) was used to organize non-parametric Wilcoxon test outputs and prepare statistical annotations for plotting.

Graphical visualization was based on the ggplot2 framework (v4.0.0) (Wickham, 2016). Extended plotting functionalities were provided by ggpubr (v0.6.1) (Kassambara, 2025) for statistical annotation layers, ggrepel (v0.9.6) (Slowikowski, 2024) for non-overlapping text labels, ggdist (v3.3.3) (Kay, 2023) for half-eye density overlays, patchwork (v1.3.2) (Pedersen, 2025) for multi-panel composition, and khroma (v1.16.0) (Frerebeau, 2025) for color-blind–friendly palettes. Additional customization relied on ggpmisc (v0.6.2) (Aphalo, 2016) and ggpp (v0.5.9) (Aphalo, 2021) for regression labels and polynomial equation annotations.

Foundational data structures and helper utilities were supported by tibble (v3.3.0) (Müller and Wickham, 2016), rlang (v1.1.6) (Henry and Wickham, 2017), and scales (v1.4.0) (Wickham et al., 2011). Ancillary visualization and formatting tools included maps (v3.4.3) (Becker et al., 2003), pheatmap (v1.0.13) (Kolde, 2010), and reshape2 (v1.4.4) (Hadley Wickham <>, 2010).

All plots were exported as vector graphics (PDF) in fixed dimensions to ensure reproducibility and consistent resolution across figures.

### Figures S1-10

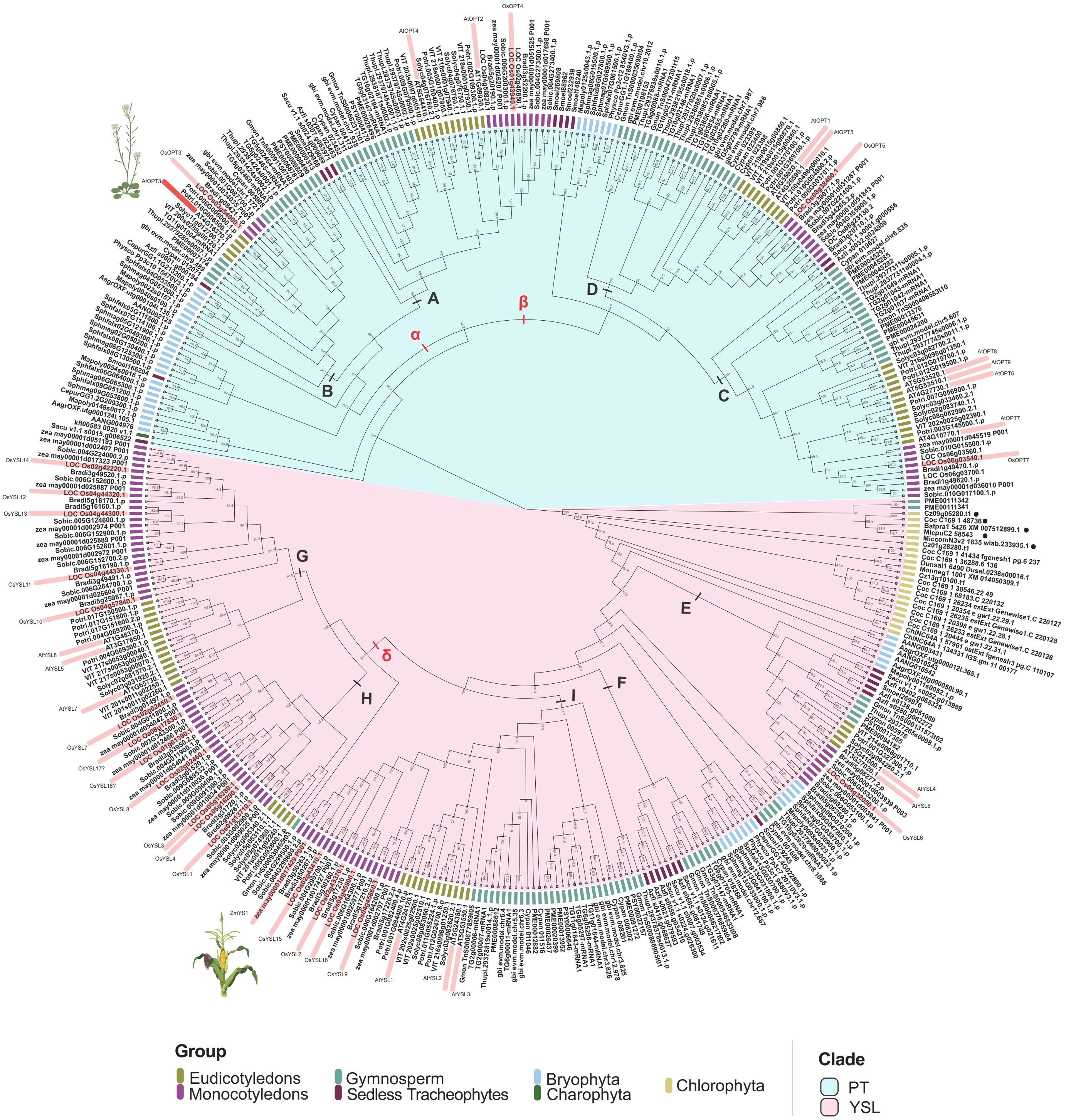


**Figure S1. Maximum-likelihood phylogenetic tree of Archaeplastida OPT proteins.** The phylogeny is shown as a cladogram with original protein identifiers and SH-like approximate likelihood-ratio test (SH-aLRT) branch support values. Selected reference proteins from *Arabidopsis thaliana*, *Oryza sativa*, and *Zea mays* discussed in the main text are highlighted in red. Branch colors indicate taxonomic groups according to the color key.

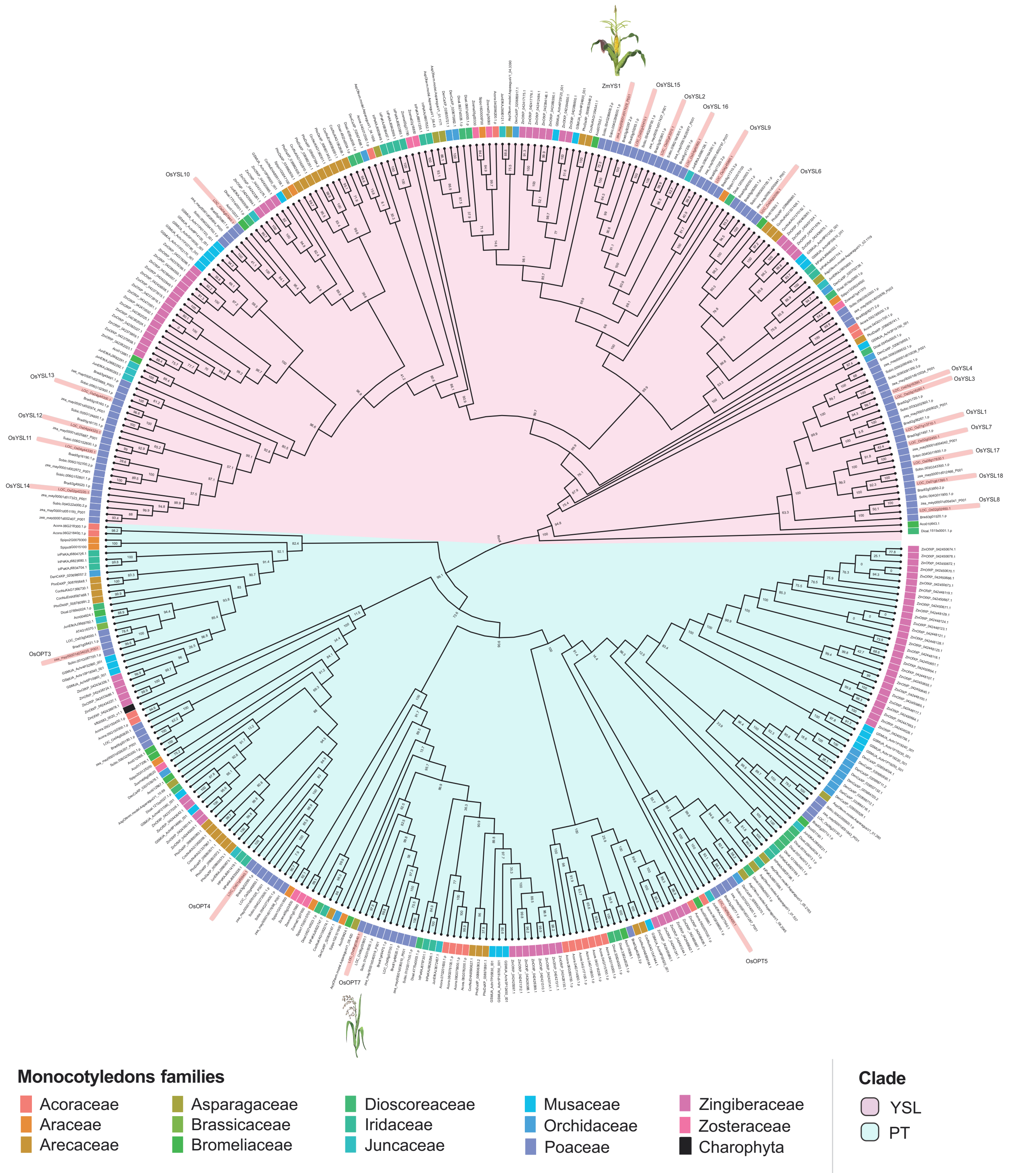


**Figure S2. Maximum-likelihood phylogenetic tree of monocot OPT proteins.** The phylogeny is shown as a cladogram with original protein identifiers and branch support values. *Oryza sativa* and *Zea mays* proteins previously defined in the Archaeplastida OPT phylogeny (Supplementary Fig. 1) are highlighted in red. *Arabidopsis thaliana* proteins are included as external references for PT and YSL subfamilies.

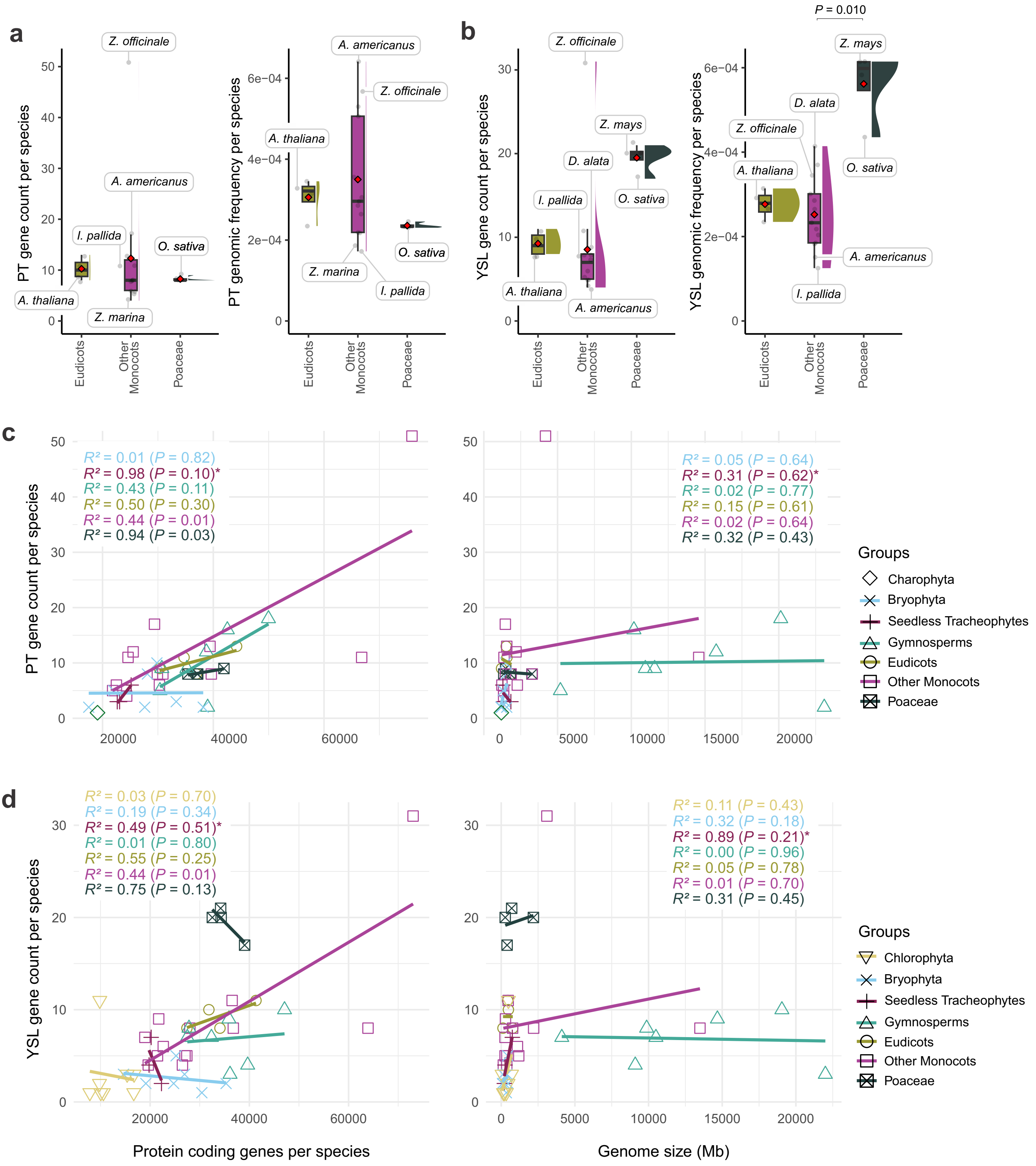


**Figure S3. YSL transporters exhibit a Poaceae-specific expansion independent of genome size.** a,b) Boxplots showing absolute gene counts (top) and normalized genomic frequencies (bottom) for PT a) and YSL b) subfamilies across eudicots, non-Poaceae monocots, and Poaceae. Central lines indicate medians; boxes span interquartile ranges; whiskers represent 1.5 x IQR; red diamonds indicate means. c,d) Correlations between PICs of PT c) or YSL (d) gene numbers and genome size or protein-coding gene number. Statistical results are provided in Tables S3 and S4.

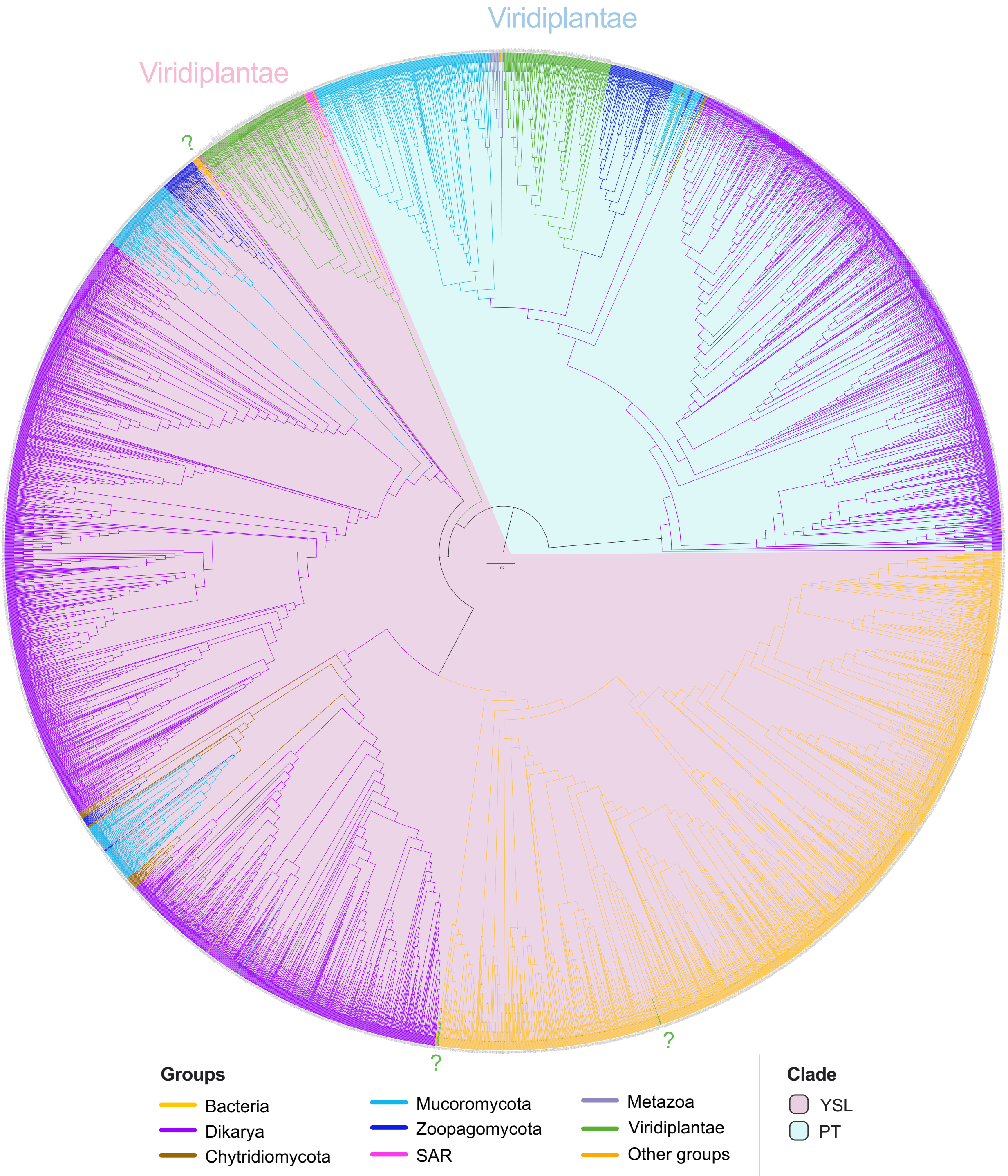


**Figure S4. Deep maximum-likelihood phylogenetic tree of OPT proteins.** The phylogeny is shown as a cladogram with original protein identifiers and SH-aLRT branch support values, highlighting the deep split between PT (blue) and YSL (pink) subfamilies. Question marks indicate chlorophyte proteins with divergent placement relative to the main Viridiplantae YSL clade. Branch colors correspond to taxonomic groups according to the color key.

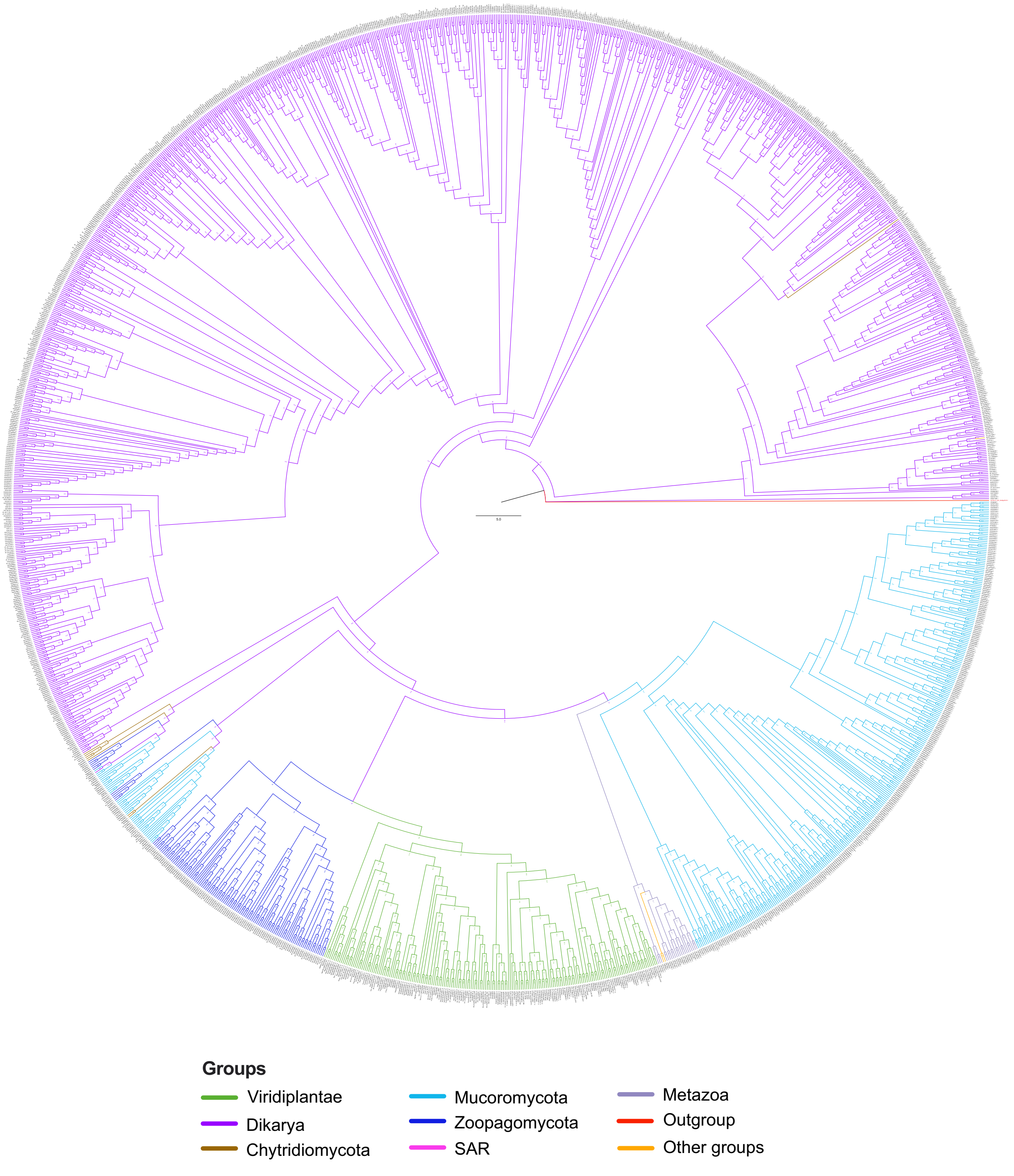


**Figure S5. Deep maximum-likelihood phylogenetic tree of the PT subfamily.** The phylogeny is shown as a cladogram with original protein identifiers and SH-aLRT branch support values and is rooted with *Oryza sativa* OsYSL15 (highlighted in red) as outgroup. Branch colors correspond to taxonomic groups according to the color key.

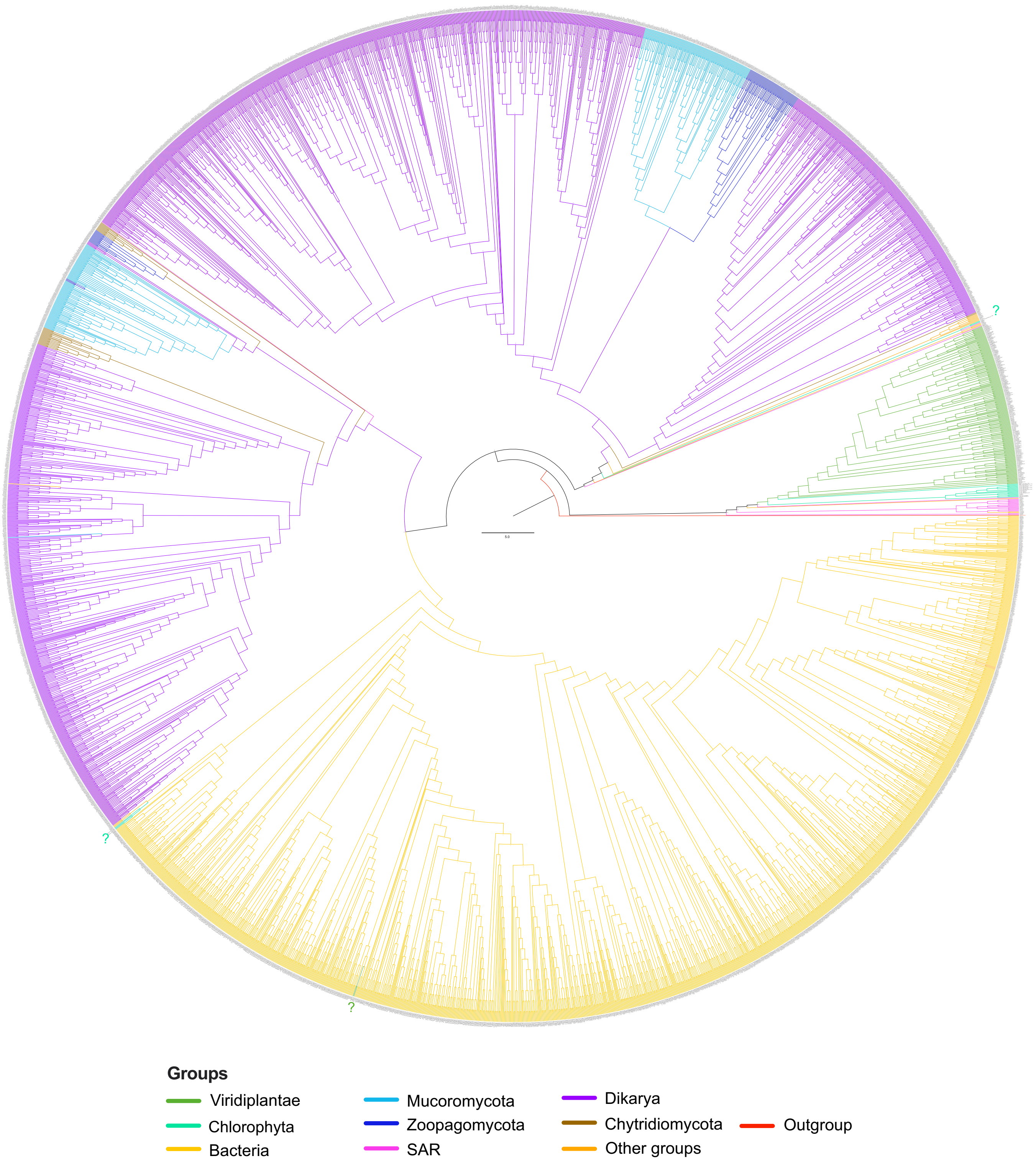


**Figure S6. Deep maximum-likelihood phylogenetic tree of the YSL subfamily.** The phylogeny is shown as a cladogram with original protein identifiers and SH-aLRT branch support values and is rooted with *Arabidopsis thaliana* AtOPT3 (highlighted in red) as outgroup. Branch colors correspond to taxonomic groups according to the color key. Question marks indicate divergent chlorophyte YSL-like proteins with fragmented phylogenetic placement.

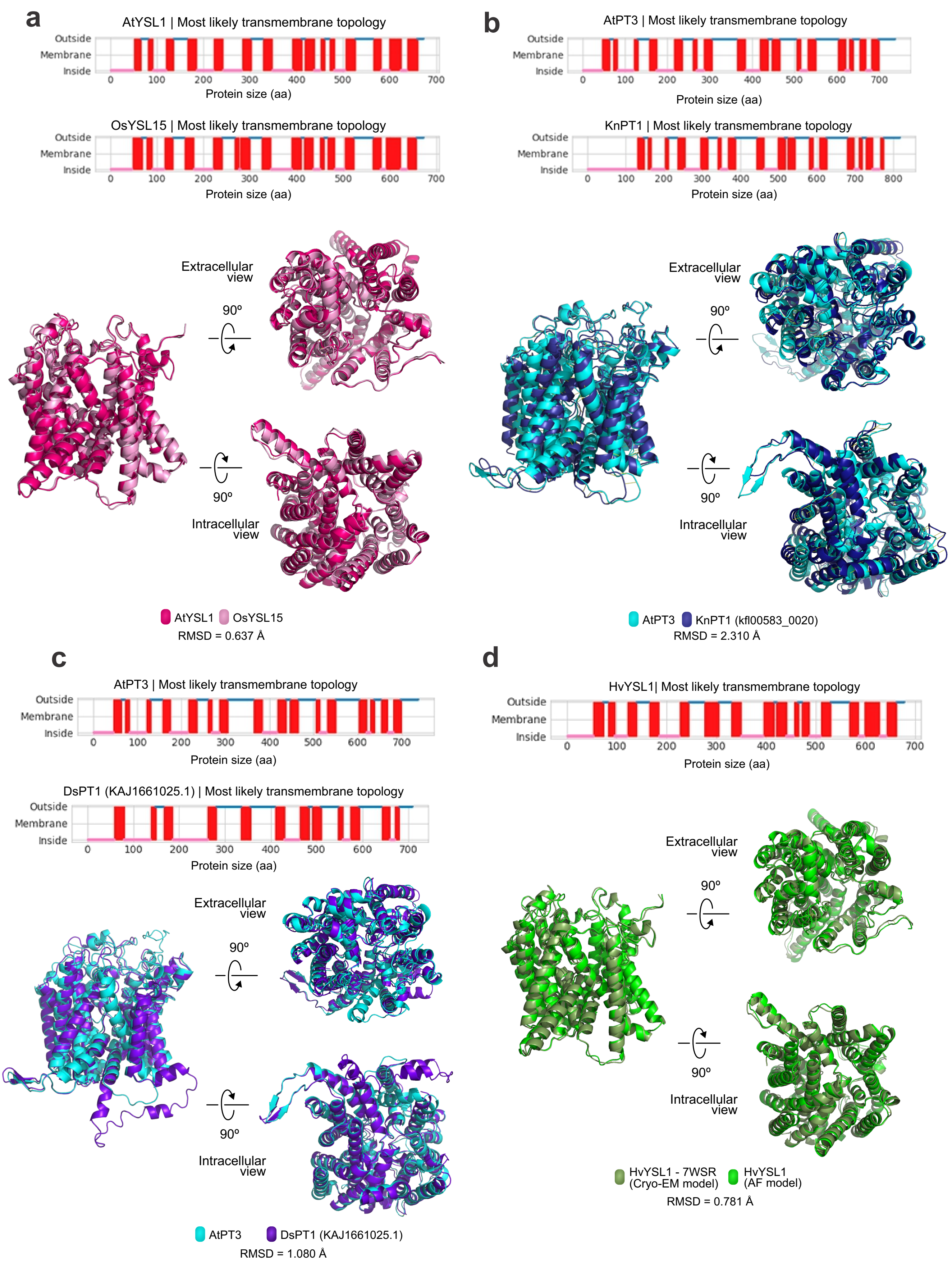


**Figure S7. Structural overview of OPT proteins across species.** a-c) Structural alignments and transmembrane topologies comparing *Arabidopsis thaliana* YSL and PT models with homologs from a) rice (*Oryza sativa*), b) *Klebsormidium nitens*, and c) the fungus *Dispira simplex*. d) Control alignment between the predicted and experimental structures of barley HvYS1. These comparisons support conserved within-subfamily architecture and structural divergence between PT and YSL proteins.

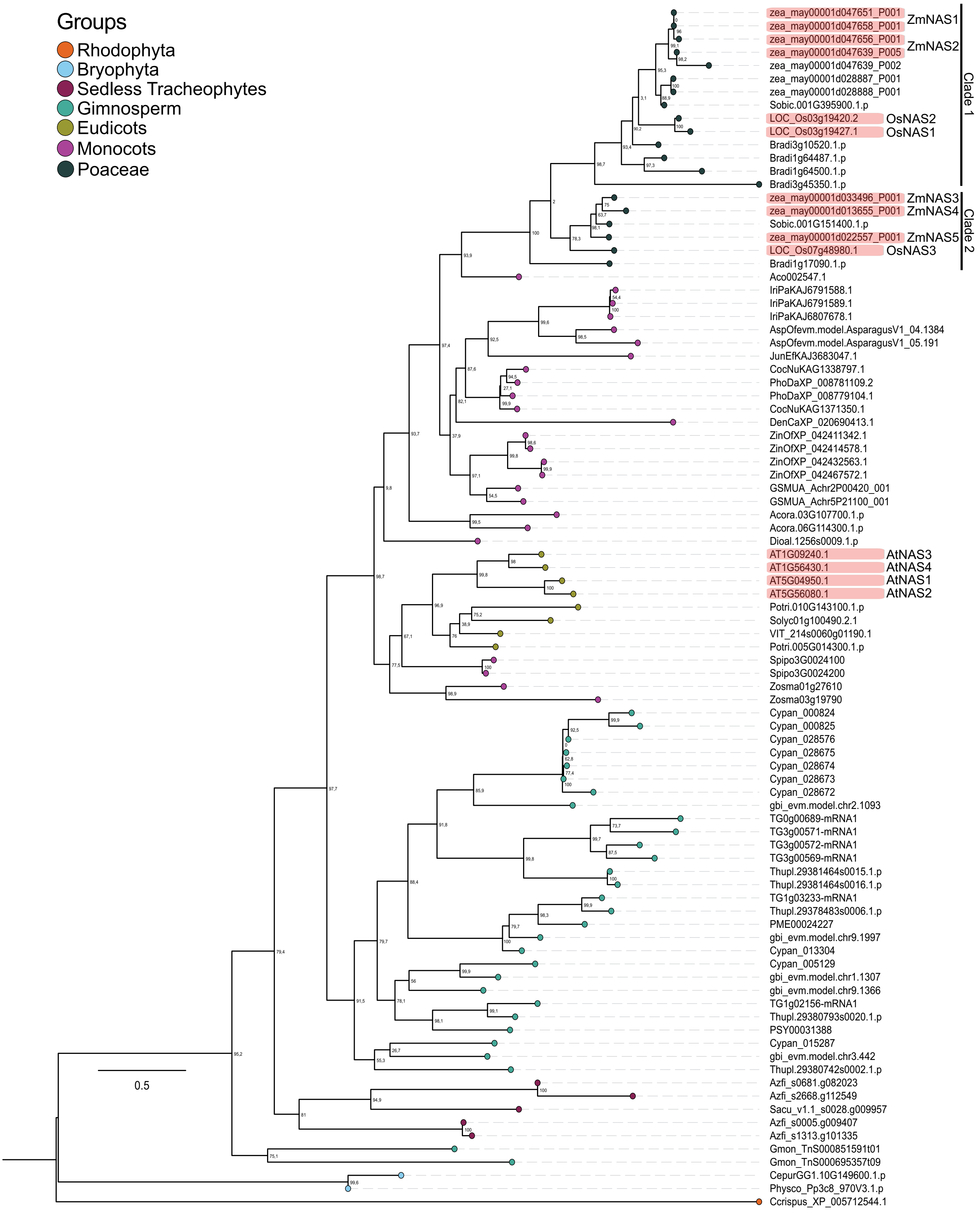


**Figure S8. Maximum-likelihood phylogenetic tree of Archaeplastida Nicotianamine Synthases (NAS).** The phylogeny is midpoint-rooted and shown as a cladogram with original protein identifiers and SH-aLRT branch support values. Branch tip colors denote taxonomic groups, and well-characterized NAS proteins from rice (*Oryza sativa*), maize (*Zea mays*), and *Arabidopsis thaliana* are highlighted in red.

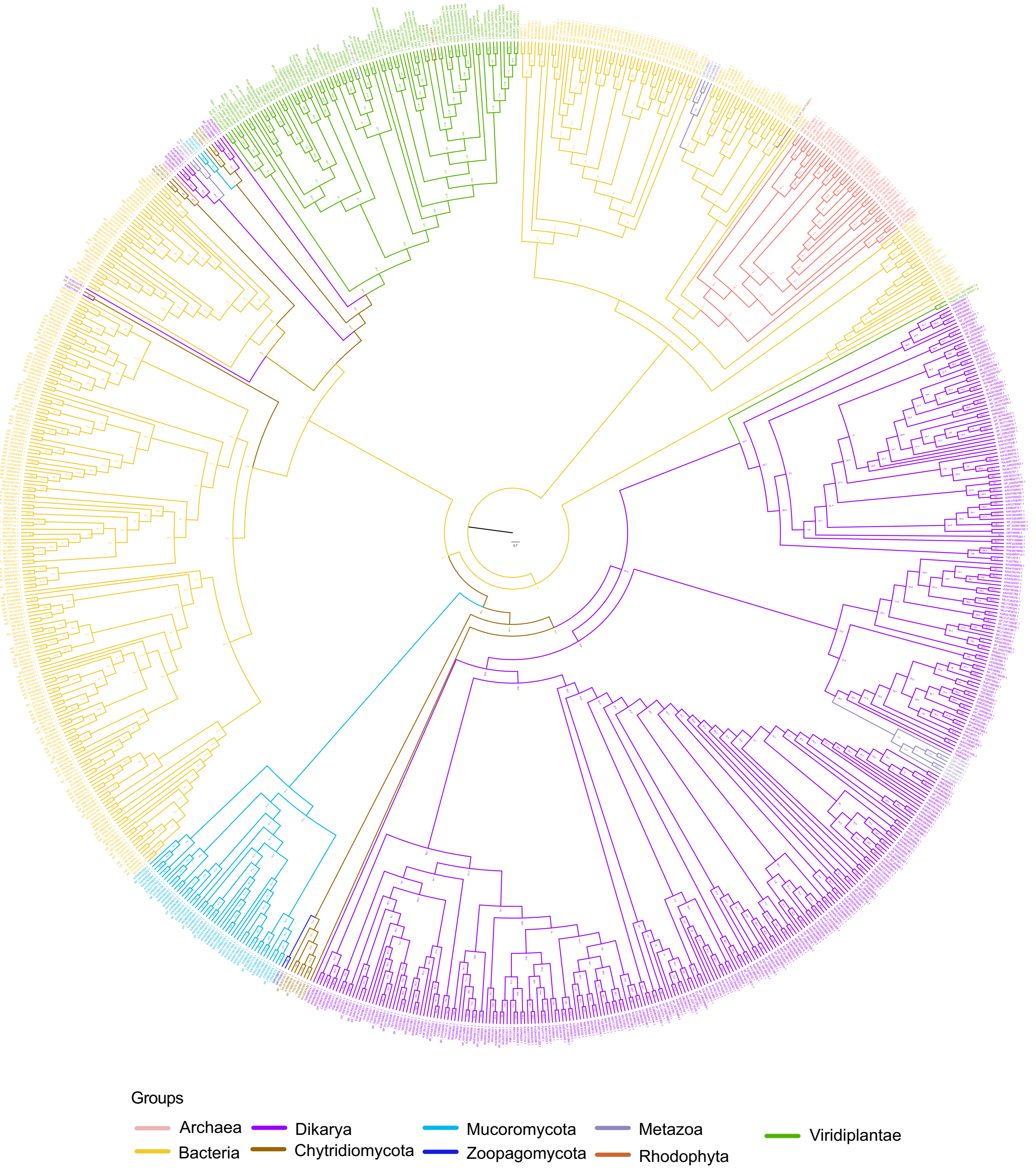


**Figure S9. Deep maximum-likelihood phylogenetic tree of Nicotianamine Synthase (NAS).** The phylogeny is midpoint-rooted and shown as a cladogram with original protein identifiers and SH-aLRT branch support values. Branch colors correspond to taxonomic groups according to the color key. Plant NAS proteins occur in separate clades associated with different fungal donors, supporting multiple independent acquisitions.

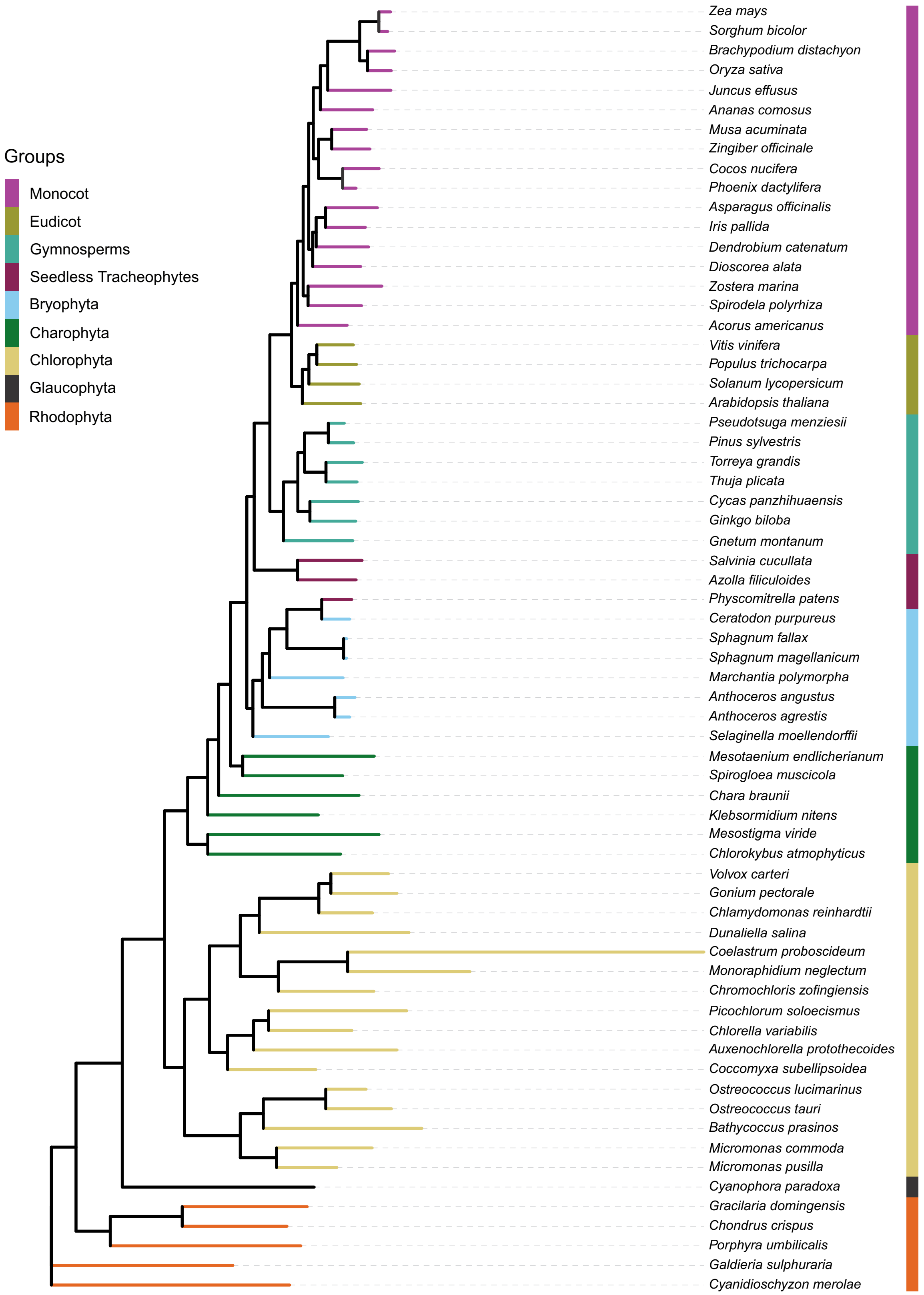


**Figure S10. Consensus species phylogeny of Archaeplastida.** This tree includes all Archaeplastida species analyzed in this study and was reconstructed from predicted proteomes. It was used for downstream analyses, including PGLS modeling and correlations based on phylogenetically independent contrasts.

### Tables S1-8

**Table S1**. Archaeplastida proteomes used in this study.

| *Species* | *Proteome Data* |
|---|---|
| *Arabidopsis thaliana* | https://phytozome-next.jgi.doe.gov/info/Athaliana_TAIR10 |
| *Populus trichocarpa* | https://phytozome-next.jgi.doe.gov/info/Ptrichocarpa_v4_1 |
| *Solanum lycopersicum* | https://phytozome-next.jgi.doe.gov/info/Slycopersicum_ITAG2_4 |
| *Vitis vinifera* | https://phytozome-next.jgi.doe.gov/info/Vvinifera_v2_1 |
| *Oryza sativa* | https://phytozome-next.jgi.doe.gov/info/Osativa_v7_0 |
| *Zea mays* | https://phytozome-next.jgi.doe.gov/info/Zmays_RefGen_V4 |
| *Sorghum bicolor* | https://phytozome-next.jgi.doe.gov/info/Sbicolor_v3_1_1 |
| *Brachypodium distachyon* | https://phytozome-next.jgi.doe.gov/info/Bdistachyon_v3_1 |
| *Acorus americanus* | https://phytozome-next.jgi.doe.gov/info/Aamericanus_v1_1 |
| *Ananas comosus* | https://phytozome-next.jgi.doe.gov/info/Acomosus_v3 |
| *Asparagus officinalis* | https://phytozome-next.jgi.doe.gov/info/Aofficinalis_V1_1 |
| *Cocos nucifera* | https://www.ncbi.nlm.nih.gov/datasets/genome/GCA_008124465.1/ |
| *Dendrobium catenatum* | https://www.ncbi.nlm.nih.gov/datasets/genome/GCF_001605985.2/ |
| *Dioscorea alata* | https://phytozome-next.jgi.doe.gov/info/Dalata_v1_1 |
| *Iris pallida* | https://www.ncbi.nlm.nih.gov/datasets/genome/GCA_029216955.1/ |
| *Juncus effusus* | https://www.ncbi.nlm.nih.gov/datasets/genome/GCA_027726005.1/ |
| *Musa acuminata* | https://phytozome-next.jgi.doe.gov/info/Macuminata_v1 |
| *Phoenix dactylifera* | https://www.ncbi.nlm.nih.gov/datasets/genome/GCF_009389715.1/ |
| *Spirodela polyrhiza* | https://phytozome-next.jgi.doe.gov/info/Spolyrhiza_v2 |
| *Zingiber officinale* | https://www.ncbi.nlm.nih.gov/datasets/genome/GCF_018446385.1/ |
| *Zostera marina* | https://phytozome-next.jgi.doe.gov/info/Zmarina_v3_1 |
| *Ginkgo biloba* | https://ngdc.cncb.ac.cn/bioproject/browse/PRJCA001755 |
| *Torreya grandis* | https://doi.org/10.6084/m9.figshare.21089869 |
| *Cycas panzhihuaensis* | https://db.cngb.org/codeplot/datasets/PwRftGHfPs5qG3gE |
| *Pinus sylvestris* | ftp://ftp.psb.ugent.be/pub/plaza/plaza_gymno_01/Fasta/proteome.psy.csv.gz |
| *Gnetum montanum* | https://datadryad.org/stash/dataset/doi:10.5061/dryad.0vm37 |
| *Pseudotsuga menziesii* | ftp://ftp.psb.ugent.be/pub/plaza/plaza_gymno_01/Fasta/proteome.pme.csv.gz |
| *Thuja plicata* | https://phytozome-next.jgi.doe.gov/info/Tplicata_v3_1 |
| *Azolla filiculoides* | https://www.fernbase.org/ftp/Azolla_filiculoides/Azolla_asm_v1.1/Azolla_filiculoides.protein.highconfidence_v1.1.fasta |
| *Salvinia cucullata* | https://www.fernbase.org/ftp/Salvinia_cucullata/Salvinia_asm_v1.2/Salvinia_cucullata.protein.highconfidence_v1.2.fasta |
| *Selaginella moellendorffii* | https://phytozome-next.jgi.doe.gov/info/Smoellendorffii_v1_0 |
| *Anthoceros agrestis* | https://www.hornworts.uzh.ch/static/download/a_agr_oxford.zip |
| *Anthoceros angustus* | https://datadryad.org/stash/dataset/doi:10.5061/dryad.msbcc2ftv |
| *Marchantia polymorpha* | https://phytozome-next.jgi.doe.gov/info/Mpolymorpha_v3_1 |
| *Sphagnum fallax* | https://phytozome-next.jgi.doe.gov/info/Sfallax_v1_1 |
| *Sphagnum magellanicum* | https://phytozome-next.jgi.doe.gov/info/Smagellanicum_v1_1 |
| *Physcomitrella patens* | https://phytozome-next.jgi.doe.gov/info/Ppatens_v3_3 |
| *Ceratodon purpureus* | https://phytozome-next.jgi.doe.gov/info/CpurpureusGG1_v1_1 |
| *Chara braunii* | https://www.ncbi.nlm.nih.gov/datasets/genome/GCA_003427395.1/ |
| *Klebsormidium nitens* | http://www.plantmorphogenesis.bio.titech.ac.jp/~algae_genome_project/klebsormidium/kf_download/160614_klebs |
| *Mesotaenium endlicherianum* | https://ndownloader.figshare.com/files/17819138 https://www.cell.com/cell/fulltext/S0092-8674(19)31169-9 |
| *Chlorokybus atmophyticus* | ftp://ftp.cngb.org/pub/CNSA/data1/CNP0000228/CNS0021447/CNA0002353/scaffold_Chlorokybus_atmophyticus.pep.gz |
| *Mesostigma viride* | ftp://ftp.cngb.org/pub/CNSA/data1/CNP0000228/CNS0021438/CNA0002352/scaffold_Mesostigma_viride.pep.gz https:/ |
| *Spirogloea muscicola* | https://ndownloader.figshare.com/files/17819147 |
| *Auxenochlorella protothecoides* | https://genome.jgi.doe.gov/portal/Auxeprot1/download/Auxeprot1_GeneCatalog_proteins_20170909.aa.fasta.gz |
| *Bathycoccus prasinos* | https://genome.jgi.doe.gov/portal/Batpra1/download/Batpra1_GeneCatalog_proteins_20180426.aa.fasta.gz |
| *Chlorella variabilis* | https://genome.jgi.doe.gov/portal/ChlNC64A_1/download/Chlorella_NC64A.all_proteins.fasta.gz |
| *Coccomyxa subellipsoidea* | https://genome.jgi.doe.gov/portal/Coc_C169_1/download/Coccomyxa_C169_v2_all_proteins.fasta.gz |
| *Chlamydomonas reinhardtii* | https://phycocosm.jgi.doe.gov/ChlreiCC4532_1/ChlreiCC4532_1.info.html |
| *Dunaliella salina* | https://genome.jgi.doe.gov/portal/Dunsal1/download/Dunsal1_GeneCatalog_proteins_20180508.aa.fasta.gz |
| *Gonium pectorale* | https://genome.jgi.doe.gov/portal/Gonpec1/download/Gonpec1_GeneCatalog_proteins_20180501.aa.fasta.gz |
| *Micromonas pusilla* | https://genome.jgi.doe.gov/portal/MicpuC2/download/MicromonasCCMP1545.allModels.aa.fasta.gz |
| *Micromonas commoda* | https://genome.jgi.doe.gov/portal/MicpuN3v2/download/MicpuN3v2_GeneCatalog_proteins_20160404.aa.fasta.gz |
| *Monoraphidium neglectum* | https://genome.jgi.doe.gov/portal/Monneg1/download/Monneg1_GeneCatalog_proteins_20170920.aa.fasta.gz |
| *Ostreococcus lucimarinus* | https://phytozome-next.jgi.doe.gov/info/Olucimarinus_v2_0 |
| *Ostreococcus tauri* | https://genome.jgi.doe.gov/portal/Ostta4221_3/download/Ostta4221_3_GeneCatalog_proteins_20161028.aa.fasta.gz |
| *Picochlorum soloecismus* | https://genome.jgi.doe.gov/portal/Picsp_1/download/Picsp_1_GeneCatalog_proteins_20170909.aa.fasta.gz |
| *Volvox carteri* | https://phytozome-next.jgi.doe.gov/info/Vcarteri_v2_1 |
| *Chromochloris zofingiensis* | https://phytozome-next.jgi.doe.gov/info/Czofingiensis_v5_2_3_2 |
| *Coelastrum proboscideum* | https://ftp.cngb.org/pub/CNSA/data2/CNP0000705/CNS0251945/CNA0014153/ |
| *Chondrus crispus* | https://phycocosm.jgi.doe.gov/Chocri1/Chocri1.home.html |
| *Cyanidioschyzon merolae* | https://phycocosm.jgi.doe.gov/Cyamer1/Cyamer1.home.html |
| *Galdieria sulphuraria* | https://phycocosm.jgi.doe.gov/Galsul1/Galsul1.home.html |
| *Porphyra umbilicalis* | https://phytozome-next.jgi.doe.gov/info/Pumbilicalis_v1_5 |
| *Gracilaria domingensis* | https://phycocosm.jgi.doe.gov/Gradom1/Gradom1.home.html |
| *Cyanophora paradoxa* | https://phycocosm.jgi.doe.gov/Cyapar1/Cyapar1.home.html |

**Table S2**. Distribution and quantitative summary of OPT and NAS homologs across Archaeplastida.

| *Species* | *Groups* | *Order* | *PCG* | *Genome Size (Mb)* | *OPT Homologues* | *GF (OPT)* | *PT Homologues* | *GF (PT)* | *YSL Homologues* | *GF (YSL)* | *NAS Homologues* | *GF (NAS)* |
|---|---|---|---|---|---|---|---|---|---|---|---|---|
| *Arabidopsis thaliana* | Eudicots | Brassicales | 27416 | 119.669 | 17 | 0.000620076 | 9 | 0.000328275 | 8 | 0.0002918 | 4 | 0.0001459 |
| *Populus trichocarpa* | Eudicots | Malpighiales | 41335 | 434.29 | 24 | 0.000580622 | 13 | 0.000314503 | 11 | 0.000266118 | 2 | 4.83851E-05 |
| *Solanum lycopersicum* | Eudicots | Malpighiales | 34075 | 782.52 | 16 | 0.000469552 | 8 | 0.000234776 | 8 | 0.000234776 | 1 | 2.9347E-05 |
| *Vitis vinifera* | Eudicots | Vitales | 31845 | 486.198 | 21 | 0.000659444 | 11 | 0.000345423 | 10 | 0.000314021 | 1 | 3.14021E-05 |
| *Oryza sativa* | Monocots | Poales | 39045 | 399.284 | 26 | 0.000665898 | 9 | 0.000230503 | 17 | 0.000435395 | 3 | 7.68344E-05 |
| *Zea mays* | Monocots | Poales | 32540 | 2182.79 | 28 | 0.000860479 | 8 | 0.000245851 | 20 | 0.000614628 | 10 | 0.000307314 |
| *Sorghum bicolor* | Monocots | Poales | 34211 | 709.345 | 29 | 0.000847681 | 8 | 0.000233843 | 21 | 0.000613838 | 2 | 5.84607E-05 |
| *Brachypodium distachyon* | Other Monocots | Poales | 34310 | 271.163 | 28 | 0.000816089 | 8 | 0.000233168 | 20 | 0.00058292 | 5 | 0.00014573 |
| *Acorus americanus* | Other Monocots | Acorales | 26508 | 375.93 | 21 | 0.000792214 | 17 | 0.000641316 | 4 | 0.000150898 | 2 | 7.54489E-05 |
| *Ananas comosus* | Other Monocots | Bromeliaceae | 27024 | 381.905 | 13 | 0.000481054 | 8 | 0.000296033 | 5 | 0.000185021 | 1 | 3.70041E-05 |
| *Asparagus officinalis* | Other Monocots | Asparagales | 27395 | 1187.539 | 11 | 0.000401533 | 6 | 0.000219018 | 5 | 0.000182515 | 2 | 7.3006E-05 |
| *Cocos nucifera* | Other Monocots | Arecales | 28016 | 2200 | 16 | 0.000571102 | 8 | 0.000285551 | 8 | 0.000285551 | 2 | 7.13878E-05 |
| *Dendrobium catenatum* | Other Monocots | Asparagales | 22643 | 1100 | 18 | 0.000794948 | 12 | 0.000529965 | 6 | 0.000264983 | 1 | 4.41638E-05 |
| *Dioscorea alata* | Other Monocots | Dioscoreales | 21728 | 287.3439 | 20 | 0.000920471 | 11 | 0.000506259 | 9 | 0.000414212 | 1 | 4.60236E-05 |
| *Iris pallida* | Other Monocots | Asparagales | 63918 | 13489.12 | 19 | 0.000297256 | 11 | 0.000172095 | 8 | 0.00012516 | 3 | 4.69351E-05 |
| *Juncus effusus* | Other Monocots | Poales | 18942 | 258.3 | 12 | 0.000633513 | 5 | 0.000263964 | 7 | 0.000369549 | 1 | 5.27927E-05 |
| *Musa acuminata* | Other Monocots | Zingiberales | 36528 | 472.9604 | 24 | 0.00065703 | 13 | 0.000355891 | 11 | 0.000301139 | 2 | 5.47525E-05 |
| *Phoenix dactylifera* | Other Monocots | Arecales | 36764 | 772.3 | 16 | 0.000435208 | 8 | 0.000217604 | 8 | 0.000217604 | 2 | 5.4401E-05 |
| *Spirodela polyrhiza* | Other Monocots | Alismatales | 19623 | 145.1994 | 10 | 0.000509606 | 6 | 0.000305764 | 4 | 0.000203842 | 2 | 0.000101921 |
| *Zingiber officinale* | Other Monocots | Zingiberales | 73003 | 3100 | 82 | 0.000912967 | 51 | 0.000567821 | 31 | 0.000345146 | 4 | 4.4535E-05 |
| *Zostera marina* | Other Monocots | Alismatales | 21483 | 260.4921 | 9 | 0.000418936 | 4 | 0.000186194 | 5 | 0.000232742 | 2 | 9.30969E-05 |
| *Ginkgo biloba* | Gymnosperms | Ginkgoales | 27832 | 9870 | 17 | 0.000610808 | 9 | 0.000323369 | 8 | 0.000287439 | 5 | 0.000179649 |
| *Torreya grandis* | Gymnosperms | Cupressales | 47089 | 19050.82 | 28 | 0.000594619 | 18 | 0.000382255 | 10 | 0.000212364 | 6 | 0.000127418 |
| *Cycas panzhihuaensis* | Gymnosperms | Cycadales | 32353 | 10500 | 16 | 0.000494545 | 9 | 0.000278181 | 7 | 0.000216363 | 10 | 0.00030909 |
| *Pinus sylvestris* | Gymnosperms | Pinales | 36106 | 22000 | 5 | 0.000138481 | 2 | 5.53925E-05 | 3 | 8.30887E-05 | 1 | 2.76962E-05 |
| *Gnetum montanum* | Gymnosperms | Gnetales | 27491 | 4110 | 12 | 0.000436506 | 5 | 0.000181878 | 7 | 0.000254629 | 2 | 7.27511E-05 |
| *Pseudotsuga menziesii* | Gymnosperms | Pinales | 35880 | 14673.2 | 21 | 0.000585284 | 12 | 0.000334448 | 9 | 0.000250836 | 1 | 2.78707E-05 |
| *Thuja plicata* | Gymnosperms | Cupressales | 39659 | 9095.91 | 20 | 0.000504299 | 16 | 0.000403439 | 4 | 0.00010086 | 5 | 0.000126075 |
| *Azolla filiculoides* | Seedless Tracheophytes | Salviniales | 20203 | 753 | 10 | 0.000494976 | 3 | 0.000148493 | 7 | 0.000346483 | 4 | 0.00019799 |
| *Salvinia cucullata* | Seedless Tracheophytes | Salviniales | 19780 | 255 | 7 | 0.000353893 | 3 | 0.000151668 | 4 | 0.000202224 | 1 | 5.05561E-05 |
| *Selaginella moellendorffii* | Seedless Tracheophytes | Selaginellales | 22285 | 212.315 | 8 | 0.000358986 | 6 | 0.000269239 | 2 | 8.97465E-05 | 0 | 0 |
| *Anthoceros agrestis* | Bryophyta | Anthocerotales | 24739 | 83 | 4 | 0.000161688 | 2 | 8.0844E-05 | 2 | 8.0844E-05 | 0 | 0 |
| *Anthoceros angustus* | Bryophyta | Anthocerotales | 14629 | 119.349 | 5 | 0.000341787 | 2 | 0.000136715 | 3 | 0.000205072 | 0 | 0 |
| *Marchantia polymorpha* | Bryophyta | Marchantiales | 19138 | 225.761 | 7 | 0.000365764 | 5 | 0.00026126 | 2 | 0.000104504 | 0 | 0 |
| *Sphagnum fallax* | Bryophyta | Sphagnales | 26939 | 395 | 13 | 0.000482572 | 10 | 0.000371209 | 3 | 0.000111363 | 0 | 0 |
| *Sphagnum magellanicum* | Bryophyta | Sphagnales | 25227 | 439.011 | 13 | 0.000515321 | 8 | 0.000317121 | 5 | 0.0001982 | 0 | 0 |
| *Physcomitrella patens* | Bryophyta | Funariales | 35307 | 472.081 | 4 | 0.000113292 | 2 | 5.6646E-05 | 2 | 5.6646E-05 | 1 | 2.8323E-05 |
| *Ceratodon purpureus* | Bryophyta | Pseudoditrichales | 30425 | 349.46 | 4 | 0.000131471 | 3 | 9.86031E-05 | 1 | 3.28677E-05 | 1 | 3.28677E-05 |
| *Chara braunii* | Charophyta | Charales | 34719 | 1751 | 0 | 0 | 0 | 0 | 0 | 0 | 0 | 0 |
| *Klebsormidium nitens* | Charophyta | Klebsormidiales | 16215 | 104.21 | 1 | 6.16713E-05 | 1 | 6.16713E-05 | 0 | 0 | 0 | 0 |
| *Mesotaenium endlicherianum* | Charophyta | Zygnematales | 11080 | 174.26 | 0 | 0 | 0 | 0 | 0 | 0 | 0 | 0 |

| | | | | | | | | | | | | |
|---|---|---|---|---|---|---|---|---|---|---|---|---|
| *Chlorokybus atmophyticus* | Charophyta | Cholokybales | 9066 | 85 | 0 | 0 | 0 | 0 | 0 | 0 | 0 | 0 |
| *Mesostigma viride* | Charophyta | Mesostigmatales | 9198 | 245 | 0 | 0 | 0 | 0 | 0 | 0 | 0 | 0 |
| *Spirogloea muscicola* | Charophyta | Spirogloeales | 27137 | 163.62 | 0 | 0 | 0 | 0 | 0 | 0 | 0 | 0 |
| *Auxenochlorella protothecoides* | Chlorophyta | Chlorellales | 7013 | 22.92 | 0 | 0 | 0 | 0 | 0 | 0 | 0 | 0 |
| *Bathycoccus prasinos* | Chlorophyta | Mamiellales | 7847 | 150.743 | 1 | 0.000127437 | 0 | 0 | 1 | 0.000127437 | 0 | 0 |
| *Chlorella variabilis* | Chlorophyta | Chlorellales | 9791 | 461.595 | 2 | 0.000204269 | 0 | 0 | 2 | 0.000204269 | 0 | 0 |
| *Coccomyxa subellipsoidea* | Chlorophyta | Trebouxiales | 9851 | 488.266 | 11 | 0.001116638 | 0 | 0 | 11 | 0.001116638 | 0 | 0 |
| *Chlamydomonas reinhardtii* | Chlorophyta | Chlamydomonadales | 16090 | 111.04 | 0 | 0 | 0 | 0 | 0 | 0 | 0 | 0 |
| *Dunaliella salina* | Chlorophyta | Chlamydomonadales | 16697 | 343.704 | 1 | 5.9891E-05 | 0 | 0 | 1 | 5.9891E-05 | 0 | 0 |
| *Gonium pectorale* | Chlorophyta | Chlamydomonadales | 16290 | 148.81 | 0 | 0 | 0 | 0 | 0 | 0 | 0 | 0 |
| *Micromonas pusilla* | Chlorophyta | Mamiellales | 10575 | 219.583 | 1 | 9.45626E-05 | 0 | 0 | 1 | 9.45626E-05 | 0 | 0 |
| *Micromonas commoda* | Chlorophyta | Mamiellales | 10056 | 211.093 | 1 | 9.94431E-05 | 0 | 0 | 1 | 9.94431E-05 | 0 | 0 |
| *Monoraphidium neglectum* | Chlorophyta | Sphaeropleales | 16761 | 697.118 | 1 | 5.96623E-05 | 0 | 0 | 3 | 0.000178987 | 0 | 0 |
| *Ostreococcus lucimarinus* | Chlorophyta | Mamiellales | 7796 | 13.205 | 0 | 0 | 0 | 0 | 0 | 0 | 0 | 0 |
| *Ostreococcus tauri* | Chlorophyta | Mamiellales | 7699 | 12.92 | 0 | 0 | 0 | 0 | 0 | 0 | 0 | 0 |
| *Picochlorum soloecismus* | Chlorophyta | Chlorellales | 6861 | 15.25 | 0 | 0 | 0 | 0 | 0 | 0 | 0 | 0 |
| *Volvox carteri* | Chlorophyta | Chlamydomonadales | 14247 | 131.163 | 0 | 0 | 0 | 0 | 0 | 0 | 0 | 0 |
| *Chromochloris zofingiensis* | Chlorophyta | Sphaeropleales | 15369 | 60.36 | 3 | 0.000195198 | 0 | 0 | 3 | 0.000195198 | 0 | 0 |
| *Coelastrum proboscideum* | Chlorophyta | Sphaeropleales | 16196 | 133.4 | 0 | 0 | 0 | 0 | 0 | 0 | 0 | 0 |
| *Chondrus crispus* | Rhodophyta | Gigartinales | 9807 | 104.8 | 0 | 0 | 0 | 0 | 0 | 0 | 1 | 0.000101968 |
| *Cyanidioschyzon merolae* | Rhodophyta | Cyanidiales | 4803 | 16.5 | 0 | 0 | 0 | 0 | 0 | 0 | 0 | 0 |
| *Galdieria sulphuraria* | Rhodophyta | Gaudieriales | 6594 | 13.7 | 0 | 0 | 0 | 0 | 0 | 0 | 0 | 0 |
| *Porphyra umbilicalis* | Rhodophyta | Bangiales | 13125 | 87.699 | 0 | 0 | 0 | 0 | 0 | 0 | 0 | 0 |
| *Gracilaria domingensis* | Rhodophyta | Gracilariales | 11532 | 77.7 | 0 | 0 | 0 | 0 | 0 | 0 | 0 | 0 |
| *Cyanophora paradoxa* | Glaucophyta | Cyanophorales | 21535 | 98.4 | 0 | 0 | 0 | 0 | 0 | 0 | 0 | 0 |

| | |
|---|---|
| *Species* | The species name analyzed in this study. |
| *Groups* | The higher-level phylogenetic group to which the species belongs. |
| *Order* | The taxonomic order of the species. |
| *PCG* | The total number of protein-coding genes annotated in the genome. |
| *Genome Size (Mb)* | The genome size in megabases (Mb). |
| *# Homologues (OPT)* | Number of identified homologs belonging to the OPT family. |
| *GF (OPT)* | Genomic frequency of OPT homologs (number of homologs normalized by total PCGs). |
| *# Homologues (PT)* | Number of identified homologs belonging to the PT family. |
| *GF (PT)* | Genomic frequency of PT homologs. |
| *# Homologues (YSL)* | Number of identified homologs belonging to the YSL family. |
| *GF (YSL)* | Genomic frequency of YSL homologs. |
| *# Homologues (NAS)* | Number of identified homologs belonging to the NAS family. |
| *GF (NAS)* | Genomic frequency of NAS homologs. |
| *Proteome Data* | Source/database link for the proteome used in the analysis. |

**Table S3**. Statistical comparisons of gene-family distributions across Archaeplastida.

| *Family/Analysis* | *Value type* | *Estimate* | *SE* | *df* | *t-ratio* | *p-value (GLS contrast)* | *group1* | *group2* | *p-value (ANOVA global)* | *p-value (Wilcoxon)* |
|---|---|---|---|---|---|---|---|---|---|---|
| OPT | Gene Count | 0.3748110 | 0.8599407 | 26 | 0.436 | 0.9994 | Chlorophyta | Charophyta | 0.0941 | 0.645387748 |
| OPT | Gene Count | -0.8708643 | 0.7406209 | 26 | -1.176 | 0.8969 | Chlorophyta | Bryophyta | 0.0941 | 0.007721469 |
| OPT | Gene Count | -1.1305695 | 0.7504273 | 26 | -1.507 | 0.7385 | Chlorophyta | Seedless Tracheophytes | 0.0941 | 0.068798413 |
| OPT | Gene Count | -1.6645191 | 0.7678314 | 26 | -2.168 | 0.3458 | Chlorophyta | Gymnosperms | 0.0941 | 0.001789843 |
| OPT | Gene Count | -1.8933425 | 0.7923695 | 26 | -2.389 | 0.2427 | Chlorophyta | Eudicots | 0.0941 | 0.006338844 |
| OPT | Gene Count | -2.2580712 | 0.8296036 | 26 | -2.722 | 0.1324 | Chlorophyta | Monocots | 0.0941 | 0.006240605 |
| OPT | Gene Count | -1.2456753 | 0.7972184 | 26 | -1.563 | 0.7059 | Charophyta | Bryophyta | 0.0941 | 0.176982150 |
| OPT | Gene Count | -1.5053804 | 0.8031835 | 26 | -1.874 | 0.5137 | Charophyta | Seedless Tracheophytes | 0.0941 | 0.500000000 |
| OPT | Gene Count | -2.0393301 | 0.8246585 | 26 | -2.473 | 0.2101 | Charophyta | Gymnosperms | 0.0941 | 0.250000000 |
| OPT | Gene Count | -2.2681535 | 0.8491133 | 26 | -2.671 | 0.1460 | Charophyta | Eudicots | 0.0941 | 0.400000000 |
| OPT | Gene Count | -2.6328822 | 0.8980520 | 26 | -2.932 | 0.0869 | Charophyta | Monocots | 0.0941 | 0.276500480 |
| OPT | Gene Count | -0.2597051 | 0.4783469 | 26 | -0.543 | 0.9978 | Bryophyta | Seedless Tracheophytes | 0.0941 | 0.416426753 |
| OPT | Gene Count | -0.7936548 | 0.5338392 | 26 | -1.487 | 0.7498 | Bryophyta | Gymnosperms | 0.0941 | 0.017332149 |
| OPT | Gene Count | -1.0224782 | 0.5722624 | 26 | -1.787 | 0.5679 | Bryophyta | Eudicots | 0.0941 | 0.009858437 |
| OPT | Gene Count | -1.3872069 | 0.6541122 | 26 | -2.121 | 0.3706 | Bryophyta | Monocots | 0.0941 | 0.009687791 |
| OPT | Gene Count | -0.5339496 | 0.5198720 | 26 | -1.027 | 0.9429 | Seedless Tracheophytes | Gymnosperms | 0.0941 | 0.116666667 |
| OPT | Gene Count | -0.7627731 | 0.5594571 | 26 | -1.363 | 0.8155 | Seedless Tracheophytes | Eudicots | 0.0941 | 0.057142857 |
| OPT | Gene Count | -1.1275018 | 0.6445999 | 26 | -1.749 | 0.5914 | Seedless Tracheophytes | Monocots | 0.0941 | 0.049745991 |
| OPT | Gene Count | -0.2288234 | 0.4931821 | 26 | -0.464 | 0.9991 | Gymnosperms | Eudicots | 0.0941 | 0.568099805 |
| OPT | Gene Count | -0.5935521 | 0.5939779 | 26 | -0.999 | 0.9496 | Gymnosperms | Monocots | 0.0941 | 0.028283862 |
| OPT | Gene Count | -0.3647287 | 0.5290669 | 26 | -0.689 | 0.9921 | Eudicots | Monocots | 0.0941 | 0.029401048 |
| OPT | Genomic Frequency | 0.0001897094 | 0.0003719112 | 26 | 0.510 | 0.9985 | Chlorophyta | Charophyta | 0.6310 | 0.666666667 |
| OPT | Genomic Frequency | -0.0000416793 | 0.0003203072 | 26 | -0.130 | 1 | Chlorophyta | Bryophyta | 0.6310 | 0.120590521 |
| OPT | Genomic Frequency | -0.0001369545 | 0.0003245483 | 26 | -0.422 | 0.9995 | Chlorophyta | Seedless Tracheophytes | 0.6310 | 0.084848485 |
| OPT | Genomic Frequency | -0.0002160763 | 0.0003320753 | 26 | -0.651 | 0.9942 | Chlorophyta | Gymnosperms | 0.6310 | 0.028904429 |
| OPT | Genomic Frequency | -0.0003180218 | 0.0003426877 | 26 | -0.928 | 0.9644 | Chlorophyta | Eudicots | 0.6310 | 0.048484848 |
| OPT | Genomic Frequency | -0.0005266167 | 0.0003587909 | 26 | -1.468 | 0.7604 | Chlorophyta | Monocots | 0.6310 | 0.048484848 |
| OPT | Genomic Frequency | -0.0002313888 | 0.0003447848 | 26 | -0.671 | 0.9931 | Charophyta | Bryophyta | 0.6310 | 0.250000000 |
| OPT | Genomic Frequency | -0.0003266640 | 0.0003473646 | 26 | -0.940 | 0.9621 | Charophyta | Seedless Tracheophytes | 0.6310 | 0.500000000 |
| OPT | Genomic Frequency | -0.0004057857 | 0.0003566522 | 26 | -1.138 | 0.9103 | Charophyta | Gymnosperms | 0.6310 | 0.250000000 |
| OPT | Genomic Frequency | -0.0005077312 | 0.0003672285 | 26 | -1.383 | 0.8058 | Charophyta | Eudicots | 0.6310 | 0.400000000 |
| OPT | Genomic Frequency | -0.0007163261 | 0.0003883938 | 26 | -1.844 | 0.5321 | Charophyta | Monocots | 0.6310 | 0.400000000 |
| OPT | Genomic Frequency | -0.0000952752 | 0.0002068777 | 26 | -0.461 | 0.9991 | Bryophyta | Seedless Tracheophytes | 0.6310 | 0.516666667 |
| OPT | Genomic Frequency | -0.0001743970 | 0.0002308773 | 26 | -0.755 | 0.9872 | Bryophyta | Gymnosperms | 0.6310 | 0.053030303 |
| OPT | Genomic Frequency | -0.0002763425 | 0.0002474947 | 26 | -1.117 | 0.9172 | Bryophyta | Eudicots | 0.6310 | 0.024242424 |
| OPT | Genomic Frequency | -0.0004849374 | 0.0002828935 | 26 | -1.714 | 0.6132 | Bryophyta | Monocots | 0.6310 | 0.006060606 |
| OPT | Genomic Frequency | -0.0000791218 | 0.0002248367 | 26 | -0.352 | 0.9998 | Seedless Tracheophytes | Gymnosperms | 0.6310 | 0.266666667 |
| OPT | Genomic Frequency | -0.0001810672 | 0.0002419567 | 26 | -0.748 | 0.9878 | Seedless Tracheophytes | Eudicots | 0.6310 | 0.114285714 |
| OPT | Genomic Frequency | -0.0003896622 | 0.0002787796 | 26 | -1.398 | 0.7981 | Seedless Tracheophytes | Monocots | 0.6310 | 0.057142857 |
| OPT | Genomic Frequency | -0.0001019455 | 0.0002132937 | 26 | -0.478 | 0.9989 | Gymnosperms | Eudicots | 0.6310 | 0.315151515 |
| OPT | Genomic Frequency | -0.0003105404 | 0.0002568864 | 26 | -1.209 | 0.8844 | Gymnosperms | Monocots | 0.6310 | 0.006060606 |
| OPT | Genomic Frequency | -0.0002085949 | 0.0002288133 | 26 | -0.912 | 0.9673 | Eudicots | Monocots | 0.6310 | 0.028571429 |
| PT | Gene Count | -0.7715592 | 10,226,958 | 19 | -0.754 | 0.9719 | Charophyta | Bryophyta | 0.6739 | 0.17971249 |
| PT | Gene Count | -0.8833626 | 10,322,446 | 19 | -0.856 | 0.9525 | Charophyta | Seedless Tracheophytes | 0.6739 | 0.34577859 |

| | | | | | | | | | | |
|---|---|---|---|---|---|---|---|---|---|---|
| PT | Gene Count | -1.4413077 | 10,540,041 | 19 | -1.367 | 0.7449 | Charophyta | Gymnosperms | 0.6739 | 0.18778127 |
| PT | Gene Count | -1.6520754 | 10,824,908 | 19 | -1.526 | 0.6527 | Charophyta | Eudicots | 0.6739 | 0.40000000 |
| PT | Gene Count | -1.4920744 | 11,265,127 | 19 | -1.325 | 0.7684 | Charophyta | Monocots | 0.6739 | 0.23567991 |
| PT | Gene Count | -0.1118033 | 0.6025918 | 19 | -0.186 | 1 | Bryophyta | Seedless Tracheophytes | 0.6739 | 0.81525212 |
| PT | Gene Count | -0.6697485 | 0.6680482 | 19 | -1.003 | 0.9114 | Bryophyta | Gymnosperms | 0.6739 | 0.06984404 |
| PT | Gene Count | -0.8805162 | 0.7147208 | 19 | -1.232 | 0.8161 | Bryophyta | Eudicots | 0.6739 | 0.03547995 |
| PT | Gene Count | -0.7205152 | 0.8017475 | 19 | -0.899 | 0.9421 | Bryophyta | Monocots | 0.6739 | 0.11819678 |
| PT | Gene Count | -0.5579451 | 0.6515409 | 19 | -0.856 | 0.9524 | Seedless Tracheophytes | Gymnosperms | 0.6739 | 0.16885536 |
| PT | Gene Count | -0.7687128 | 0.6996138 | 19 | -1.099 | 0.8759 | Seedless Tracheophytes | Eudicots | 0.6739 | 0.04974599 |
| PT | Gene Count | -0.6087119 | 0.7908022 | 19 | -0.770 | 0.9694 | Seedless Tracheophytes | Monocots | 0.6739 | 0.04158634 |
| PT | Gene Count | -0.2107677 | 0.6147652 | 19 | -0.343 | 0.9993 | Gymnosperms | Eudicots | 0.6739 | 1 |
| PT | Gene Count | -0.0507667 | 0.7272627 | 19 | -0.070 | 1 | Gymnosperms | Monocots | 0.6739 | 0.38631627 |
| PT | Gene Count | 0.1600010 | 0.6476847 | 19 | 0.247 | 0.9999 | Eudicots | Monocots | 0.6739 | 0.16347531 |
| PT | Genomic Frequency | -1.081498 | 0.0002140083 | 19 | -0.505 | 0.9953 | Charophyta | Bryophyta | 0.9161 | 0.50000000 |
| PT | Genomic Frequency | -1.299747 | 0.0002160065 | 19 | -0.602 | 0.9896 | Charophyta | Seedless Tracheophytes | 0.9161 | 0.50000000 |
| PT | Genomic Frequency | -1.934109 | 0.0002205599 | 19 | -0.877 | 0.9475 | Charophyta | Gymnosperms | 0.9161 | 0.50000000 |
| PT | Genomic Frequency | -2.351785 | 0.0002265210 | 19 | -1.038 | 0.8990 | Charophyta | Eudicots | 0.9161 | 0.40000000 |
| PT | Genomic Frequency | -1.657343 | 0.0002357329 | 19 | -0.703 | 0.9793 | Charophyta | Monocots | 0.9161 | 0.40000000 |
| PT | Genomic Frequency | -2.182493 | 0.0001260978 | 19 | -0.173 | 1 | Bryophyta | Seedless Tracheophytes | 0.9161 | 0.66666667 |
| PT | Genomic Frequency | -8.526113 | 0.0001397951 | 19 | -0.610 | 0.9890 | Bryophyta | Gymnosperms | 0.9161 | 0.20862471 |
| PT | Genomic Frequency | -1.270287 | 0.0001495618 | 19 | -0.849 | 0.9539 | Bryophyta | Eudicots | 0.9161 | 0.23030303 |
| PT | Genomic Frequency | -5.758452 | 0.0001677729 | 19 | -0.343 | 0.9993 | Bryophyta | Monocots | 0.9161 | 0.78787879 |
| PT | Genomic Frequency | -6.34362 | 0.0001363408 | 19 | -0.465 | 0.9968 | Seedless Tracheophytes | Gymnosperms | 0.9161 | 0.18333333 |
| PT | Genomic Frequency | -1.052038 | 0.0001464005 | 19 | -0.719 | 0.9772 | Seedless Tracheophytes | Eudicots | 0.9161 | 0.11428571 |
| PT | Genomic Frequency | -3.575959 | 0.0001654825 | 19 | -0.216 | 0.9999 | Seedless Tracheophytes | Monocots | 0.9161 | 0.62857143 |
| PT | Genomic Frequency | -4.176759 | 0.0001286452 | 19 | -0.325 | 0.9994 | Gymnosperms | Eudicots | 0.9161 | 1 |
| PT | Genomic Frequency | 2.767661 | 0.0001521863 | 19 | 0.182 | 1 | Gymnosperms | Monocots | 0.9161 | 0.31515152 |
| PT | Genomic Frequency | 6.944419 | 0.0001355339 | 19 | 0.512 | 0.9950 | Eudicots | Monocots | 0.9161 | 0.05714286 |
| YSL | Gene Count | -0.0943729 | 0.7825558 | 26 | -0.121 | 1 | Chlorophyta | Bryophyta | 0.2192 | 0.471169998 |
| YSL | Gene Count | -0.3812151 | 0.7929175 | 26 | -0.481 | 0.9965 | Chlorophyta | Seedless Tracheophytes | 0.2192 | 0.207821389 |
| YSL | Gene Count | -0.8719973 | 0.8113070 | 26 | -1.075 | 0.8869 | Chlorophyta | Gymnosperms | 0.2192 | 0.022164847 |
| YSL | Gene Count | -1.1430452 | 0.8372345 | 26 | -1.365 | 0.7465 | Chlorophyta | Eudicots | 0.2192 | 0.036973144 |
| YSL | Gene Count | -1.8501531 | 0.8765768 | 26 | -2.111 | 0.3130 | Chlorophyta | Monocots | 0.2192 | 0.007154342 |
| YSL | Gene Count | -0.2868422 | 0.5054315 | 26 | -0.568 | 0.9923 | Bryophyta | Seedless Tracheophytes | 0.2192 | 0.288397499 |
| YSL | Gene Count | -0.7776244 | 0.5640658 | 26 | -1.379 | 0.7389 | Bryophyta | Gymnosperms | 0.2192 | 0.006723863 |
| YSL | Gene Count | -1.0486723 | 0.6046646 | 26 | -1.734 | 0.5228 | Bryophyta | Eudicots | 0.2192 | 0.009687791 |
| YSL | Gene Count | -1.7557802 | 0.6911489 | 26 | -2.54 | 0.1491 | Bryophyta | Monocots | 0.2192 | 0.009687791 |
| YSL | Gene Count | -0.4907822 | 0.5493078 | 26 | -0.893 | 0.9447 | Seedless Tracheophytes | Gymnosperms | 0.2192 | 0.203017106 |
| YSL | Gene Count | -0.7618301 | 0.5911343 | 26 | -1.289 | 0.7883 | Seedless Tracheophytes | Eudicots | 0.2192 | 0.049745991 |
| YSL | Gene Count | -1.468938 | 0.6810980 | 26 | -2.157 | 0.2912 | Seedless Tracheophytes | Monocots | 0.2192 | 0.049745991 |
| YSL | Gene Count | -0.2710479 | 0.5211068 | 26 | -0.520 | 0.9949 | Gymnosperms | Eudicots | 0.2192 | 0.125299103 |
| YSL | Gene Count | -0.9781558 | 0.6276097 | 26 | -1.559 | 0.6314 | Gymnosperms | Monocots | 0.2192 | 0.010379103 |
| YSL | Gene Count | -0.7071079 | 0.5590233 | 26 | -1.265 | 0.8007 | Eudicots | Monocots | 0.2192 | 0.028429536 |
| YSL | Genomic Frequency | 0.0001367595 | 0.0002545425 | 26 | 0.537 | 0.9940 | Chlorophyta | Bryophyta | 0.5458 | 0.396891997 |
| YSL | Genomic Frequency | 0.0000645081 | 0.0002579128 | 26 | 0.250 | 0.9998 | Chlorophyta | Seedless Tracheophytes | 0.5458 | 0.775757576 |
| YSL | Genomic Frequency | 0.0000488553 | 0.0002638944 | 26 | 0.185 | 1 | Chlorophyta | Gymnosperms | 0.5458 | 0.231857032 |
| YSL | Genomic Frequency | -0.0000121341 | 0.0002723278 | 26 | -0.045 | 1 | Chlorophyta | Eudicots | 0.5458 | 0.048484848 |
| YSL | Genomic Frequency | -0.0002901800 | 0.0002851247 | 26 | -1.018 | 0.9078 | Chlorophyta | Monocots | 0.5458 | 0.048484848 |

| | | | | | | | | | | |
|---|---|---|---|---|---|---|---|---|---|---|
| YSL | Genomic Frequency | -0.0000722514 | 0.0001644020 | 26 | -0.439 | 0.9977 | Bryophyta | Seedless Tracheophytes | 0.5458 | 0.266666667 |
| YSL | Genomic Frequency | -0.0000879041 | 0.0001834741 | 26 | -0.479 | 0.9965 | Bryophyta | Gymnosperms | 0.5458 | 0.037878788 |
| YSL | Genomic Frequency | -0.0001488936 | 0.0001966797 | 26 | -0.757 | 0.9723 | Bryophyta | Eudicots | 0.5458 | 0.006060606 |
| YSL | Genomic Frequency | -0.0004269395 | 0.0002248105 | 26 | -1.899 | 0.4249 | Bryophyta | Monocots | 0.5458 | 0.006060606 |
| YSL | Genomic Frequency | -0.0000156528 | 0.0001786737 | 26 | -0.088 | 1 | Seedless Tracheophytes | Gymnosperms | 0.5458 | 1 |
| YSL | Genomic Frequency | -0.0000766422 | 0.0001922787 | 26 | -0.399 | 0.9985 | Seedless Tracheophytes | Eudicots | 0.5458 | 0.628571429 |
| YSL | Genomic Frequency | -0.0003546881 | 0.0002215412 | 26 | -1.601 | 0.6052 | Seedless Tracheophytes | Monocots | 0.5458 | 0.057142857 |
| YSL | Genomic Frequency | -0.0000609894 | 0.0001695008 | 26 | -0.360 | 0.9991 | Gymnosperms | Eudicots | 0.5458 | 0.072727273 |
| YSL | Genomic Frequency | -0.0003390354 | 0.0002041430 | 26 | -1.661 | 0.5681 | Gymnosperms | Monocots | 0.5458 | 0.006060606 |
| YSL | Genomic Frequency | -0.0002780459 | 0.0001818339 | 26 | -1.529 | 0.6495 | Eudicots | Monocots | 0.5458 | 0.028571429 |
| YSL | Gene Count | 0.3254691 | 0.3730220 | 16 | 0.8725198 | 0.6647 | Eudicotyledons | Other Monocots | 0.0524 | 0.13465725 |
| YSL | Gene Count | -0.5845174 | 0.4898946 | 16 | -1.1931493 | 0.4739 | Eudicotyledons | Poaceae | 0.0524 | 0.02842954 |
| YSL | Gene Count | -0.9099865 | 0.3531412 | 16 | -2.5768345 | 0.0503 | Other Monocots | Poaceae | 0.0524 | 0.01430743 |
| YSL | Genomic Frequency | 3.347244 | 8.723276 | 16 | 0.3837142 | 0.9224 | Eudicotyledons | Other Monocots | 0.0128 | 0.41176470 |
| YSL | Genomic Frequency | -2.451794 | 1.145639 | 16 | -2.1401102 | 0.1130 | Eudicotyledons | Poaceae | 0.0128 | 0.02857142 |
| YSL | Genomic Frequency | -2.786518 | 8.258356 | 16 | -3.3741805 | 0.0102 | Other Monocots | Poaceae | 0.0128 | 0.00084033 |
| NAS | Gene Count | -0.07967557 | 1.537098 | 25 | -0.05183505 | 1 | Rhodophyta | Bryopsyda | 0.8673 | NA |
| NAS | Gene Count | -0.55321372 | 1.504095 | 25 | -0.36780499 | 0.9997 | Rhodophyta | Salviniales | 0.8673 | 1 |
| NAS | Gene Count | -0.85281875 | 1.45704 | 25 | -0.58530920 | 0.9967 | Rhodophyta | Gymnosperms | 0.8673 | 0.36808283 |
| NAS | Gene Count | -0.47282999 | 1.483888 | 25 | -0.31864257 | 0.9998 | Rhodophyta | Eudicots | 0.8673 | 0.69263278 |
| NAS | Gene Count | -0.45395893 | 1.451573 | 25 | -0.31273584 | 0.9999 | Rhodophyta | Other monocots | 0.8673 | 0.27693742 |
| NAS | Gene Count | -1.15981331 | 1.523679 | 25 | -0.76119270 | 0.9866 | Rhodophyta | Poaceae | 0.8673 | 0.40000000 |
| NAS | Gene Count | -0.47353816 | 1.053223 | 25 | -0.44960846 | 0.9992 | Bryopsyda | Salviniales | 0.8673 | 0.61707508 |
| NAS | Gene Count | -0.77314318 | 0.9807888 | 25 | -0.78828709 | 0.9840 | Bryopsyda | Gymnosperms | 0.8673 | 0.16686575 |
| NAS | Gene Count | -0.39315443 | 1.026553 | 25 | -0.38298510 | 0.9996 | Bryopsyda | Eudicots | 0.8673 | 0.41131379 |
| NAS | Gene Count | -0.37428336 | 0.9891706 | 25 | -0.37838102 | 0.9997 | Bryopsyda | Other monocots | 0.8673 | 0.11467960 |
| NAS | Gene Count | -1.08013774 | 1.111721 | 25 | -0.97159035 | 0.9556 | Bryopsyda | Poaceae | 0.8673 | 0.10020951 |
| NAS | Gene Count | -0.29960502 | 0.8902349 | 25 | -0.33654603 | 0.9998 | Salviniales | Gymnosperms | 0.8673 | 0.45466093 |
| NAS | Gene Count | 0.08038373 | 0.9408652 | 25 | 0.08543597 | 1 | Salviniales | Eudicots | 0.8673 | 1 |
| NAS | Gene Count | 0.09925480 | 0.9006403 | 25 | 0.11020471 | 0.9999 | Salviniales | Other monocots | 0.8673 | 0.85489277 |
| NAS | Gene Count | -0.60659958 | 1.0351 | 25 | -0.58603022 | 0.9966 | Salviniales | Poaceae | 0.8673 | 0.53333333 |
| NAS | Gene Count | 0.37998876 | 0.7326885 | 25 | 0.51862251 | 0.9983 | Gymnosperms | Eudicots | 0.8673 | 0.24355653 |
| NAS | Gene Count | 0.39885982 | 0.6831354 | 25 | 0.58386642 | 0.9967 | Gymnosperms | Other monocots | 0.8673 | 0.14597689 |
| NAS | Gene Count | -0.30699456 | 0.8576540 | 25 | -0.35794686 | 0.9997 | Gymnosperms | Poaceae | 0.8673 | 0.70155190 |
| NAS | Gene Count | 0.01887107 | 0.5398321 | 25 | 0.03495728 | 1 | Eudicots | Other monocots | 0.8673 | 0.85409101 |
| NAS | Gene Count | -0.68698331 | 0.7556422 | 25 | -0.90913836 | 0.9676 | Eudicots | Poaceae | 0.8673 | 0.14405063 |
| NAS | Gene Count | -0.70585438 | 0.5653039 | 25 | -1.24862827 | 0.8680 | Other monocots | Poaceae | 0.8673 | 0.02585087 |
| NAS | Genomic Frequency | 5.757355 | 2.570228 | 25 | 0.224001705 | 0.9999 | Rhodophyta | Bryopsyda | 0.9722 | 0.66666667 |
| NAS | Genomic Frequency | -3.861871 | 2.515043 | 25 | -0.153550885 | 0.9999 | Rhodophyta | Salviniales | 0.9722 | 1 |
| NAS | Genomic Frequency | -3.234032 | 2.43636 | 25 | -0.132740292 | 0.9999 | Rhodophyta | Gymnosperms | 0.9722 | 1 |
| NAS | Genomic Frequency | 1.316322 | 2.481255 | 25 | 0.053050677 | 1 | Rhodophyta | Eudicots | 0.9722 | 0.80000000 |
| NAS | Genomic Frequency | 1.356581 | 2.427219 | 25 | 0.055890347 | 1 | Rhodophyta | Other monocots | 0.9722 | 0.14285714 |
| NAS | Genomic Frequency | -6.9199 | 2.54779 | 25 | -0.271604043 | 0.9999 | Rhodophyta | Poaceae | 0.9722 | 1 |
| NAS | Genomic Frequency | -9.619226 | 1.761127 | 25 | -0.546197270 | 0.9977 | Bryopsyda | Salviniales | 0.9722 | 0.33333333 |
| NAS | Genomic Frequency | -8.991387 | 1.640007 | 25 | -0.548253068 | 0.9977 | Bryopsyda | Gymnosperms | 0.9722 | 0.50000000 |
| NAS | Genomic Frequency | -4.441033 | 1.71653 | 25 | -0.258721548 | 0.9999 | Bryopsyda | Eudicots | 0.9722 | 0.53333333 |
| NAS | Genomic Frequency | -4.400774 | 1.654022 | 25 | -0.266065025 | 0.9999 | Bryopsyda | Other monocots | 0.9722 | 0.01904762 |
| NAS | Genomic Frequency | -1.267725 | 1.858943 | 25 | -0.681960442 | 0.9924 | Bryopsyda | Poaceae | 0.9722 | 0.13333333 |

| | | | | | | | | | | |
|---|---|---|---|---|---|---|---|---|---|---|
| NAS | Genomic Frequency | 6.278393 | 1.488589 | 25 | 0.042176813 | 1 | Salviniales | Gymnosperms | 0.9722 | 0.88888889 |
| NAS | Genomic Frequency | 5.178193 | 1.573249 | 25 | 0.329140051 | 0.9998 | Salviniales | Eudicots | 0.9722 | 0.26666667 |
| NAS | Genomic Frequency | 5.218452 | 1.505988 | 25 | 0.346513572 | 0.9998 | Salviniales | Other monocots | 0.9722 | 0.47619048 |
| NAS | Genomic Frequency | -3.058029 | 1.730821 | 25 | -0.176680814 | 0.9999 | Salviniales | Poaceae | 0.9722 | 0.80000000 |
| NAS | Genomic Frequency | 4.550354 | 1.225151 | 25 | 0.371411795 | 0.9997 | Gymnosperms | Eudicots | 0.9722 | 0.64848485 |
| NAS | Genomic Frequency | 4.590613 | 1.142291 | 25 | 0.401877582 | 0.9996 | Gymnosperms | Other monocots | 0.9722 | 0.24135707 |
| NAS | Genomic Frequency | -3.685868 | 1.434109 | 25 | -0.257014475 | 0.9999 | Gymnosperms | Poaceae | 0.9722 | 0.64848485 |
| NAS | Genomic Frequency | 0.4025893 | 9.026696 | 25 | 0.004459986 | 1 | Eudicots | Other monocots | 0.9722 | 0.41176471 |
| NAS | Genomic Frequency | -8.236222 | 1.263532 | 25 | -0.651841105 | 0.9940 | Eudicots | Poaceae | 0.9722 | 0.20000000 |
| NAS | Genomic Frequency | -8.276481 | 9.452617 | 25 | -0.875575650 | 0.9730 | Other monocots | Poaceae | 0.9722 | 0.03193277 |

| | |
|---|---|
| *Family/Analysis* | Gene family analyzed (e.g., OPT) and type of data (Gene Count or Genomic Frequency). |
| *Value type* | Variable being tested (Gene Count or Genomic Frequency). |
| *Estimate* | GLS estimate for the difference between groups. |
| *SE* | Standard error of the GLS estimate. |
| *df* | Degrees of freedom in the GLS model. |
| *t.ratio* | t-statistic from GLS. |
| *p.value (GLS contrast)* | p-value of the GLS contrast between group1 and group2. |
| *group1* | First clade being compared in the contrast. |
| *group2* | Second clade being compared in the contrast. |
| *p-value (ANOVA global)* | p-value of the global ANOVA under the GLS (Brownian motion) model. |
| *p-value (Wilcoxon)* | p-value of the non-phylogenetic Wilcoxon test (reported separately). |

**Table S4**. Correlation and linear regression analyses based on Phylogenetically Independent Contrasts (PICs).

| ***PT*** | | | ***gene count X protein coding genes*** | | | | | | ***gene count X genome size*** | | | | | |
|---|---|---|---|---|---|---|---|---|---|---|---|---|---|---|
| *Group* | *n_tips* | *n_pic* | *r2_cor* | *p_cor* | *r2_lm0* | *p_lm0* | *r2_simple* | *p_simple* | *r2_cor* | *p_cor* | *r2_lm0* | *p_lm0* | *r2_simple* | *p_simple* |
| Charophyta | 1 | 0 | NA | NA | NA | NA | NA | NA | NA | NA | NA | NA | NA | NA |
| Bryophyta | 7 | 6 | 0.00296 | 0.919 | 0.0115 | 0.819 | 0.01 | 0.986 | 0.00132 | 0.946 | 0.0466 | 0.642 | 0.23 | 0.275 |
| Seedless Tracheophytes | 3 | 2 | NA | NA | NA | NA | 0.98 | 0.101 | NA | NA | NA | NA | 0.31 | 0.621 |
| Gymnosperms | 7 | 6 | 0.513 | 0.109 | 0.428 | 0.111 | 0.49 | 0.080 | 0.000239 | 0.977 | 0.0188 | 0.770 | 0.01 | 0.947 |
| Monocots | 13 | 12 | 0.461 | 0.151 | 0.444 | 0.012 | 0.55 | 0.043 | 0.022 | 0.646 | 0.02 | 0.064 | 0.02 | 0.635 |
| Eudicots | 4 | 3 | 0.468 | 0.520 | 0.499 | 0.293 | 0.47 | 0.312 | 0.0603 | 0.842 | 0.146 | 0.618 | 0.01 | 0.805 |
| Poaceae | 4 | 3 | 0.911 | 0.192 | 0.944 | 0.0283 | 0.92 | 0.043 | 0.655 | 0.400 | 0.325 | 0.430 | 0.14 | 0.628 |
| ***YSL*** | | | ***gene count X protein coding genes*** | | | | | | ***gene count X genome size*** | | | | | |
| *Group* | *n_tips* | *n_pic* | *r2_cor* | *p_cor* | *r2_lm0* | *p_lm0* | *r2_simple* | *p_simple* | *r2_cor* | *p_cor* | *r2_lm0* | *p_lm0* | *r2_simple* | *p_simple* |
| Chlorophyta | 8 | 7 | 0.000631 | 0.957 | 0.0260 | 0.703 | 0.01 | 0.797 | 0.0881 | 0.518 | 0.106 | 0.432 | 0.15 | 0.346 |
| Bryophyta | 7 | 6 | 0.161 | 0.430 | 0.181 | 0.341 | 0.07 | 0.562 | 0.250 | 0.313 | 0.323 | 0.183 | 0.06 | 0.597 |
| Seedless Tracheophytes | 3 | 2 | NA | NA | NA | NA | 0.49 | 0.507 | NA | NA | NA | NA | 0.89 | 0.215 |
| Gymnosperms | 7 | 6 | 0.0388 | 0.708 | 0.0141 | 0.800 | 0.01 | 0.797 | 0.168 | 0.420 | 0.000670 | 0.956 | 0.01 | 0.890 |
| Eudicots | 4 | 3 | 0.547 | 0.470 | 0.555 | 0.255 | 0.55 | 0.260 | 0.0285 | 0.892 | 0.0493 | 0.778 | 0.01 | 0.997 |
| Monocots | 13 | 12 | 0.435 | 0.0196 | 0.436 | 0.0140 | 0.59 | 0.002 | 0.0113 | 0.742 | 0.0142 | 0.698 | 0.03 | 0.593 |
| Poaceae | 4 | 3 | 0.674 | 0.387 | 0.751 | 0.133 | 0.78 | 0.115 | 0.354 | 0.594 | 0.306 | 0.447 | 0.08 | 0.718 |

| | |
|---|---|
| *Group* | The clade/group of species where the calculation was performed. |
| *n_tips* | Number of species from that clade found in the phylogenetic tree. |
| *n_pic* | Number of independent contrasts (PICs) actually used. |
| *r2_cor* | The $R^2$ of the Pearson correlation between PICs of gene count and protein-coding genes or genome size. |
| *p_cor* | The p-value of the Pearson correlation test between PICs. |
| *r2_lm0* | The $R^2$ of the linear regression without intercept (forced through the origin) on PICs. |
| *p_lm0* | The p-value of the regression coefficient (slope) in the no-intercept model. |
| *r2_simple* | The $R^2$ of the simple linear regression (non-phylogenetic). |
| *p_simple* | The p-value of the simple linear regression. |

**Table S5.** Identification of Nicotianamine Synthase (NAS) homologs in bryophyte and lycophyte transcriptomes from the 1,000 Plants (1KP) project.

| *Group* | *Subgroup* | *Family* | *Species* | *Code (OneKP)* | *# Homologues* |
|---|---|---|---|---|---|
| Bryophyta | Hornworts | Notothyladaceae | *Anthoceros agrestisB* | BSNI | NA |
| Bryophyta | Hornworts | Notothyladaceae | *Anthoceros agrestisA* | TWUW | NA |
| Bryophyta | Hornworts | Anthoceroceae | *Anthoceros formosae* | IQJU | NA |
| Bryophyta | Hornworts | Anthoceroceae | *Paraphymatoceros hallii* | FAJB | NA |
| Bryophyta | Hornworts | Dendrocerotaceae | *Phaeomegaceros coriaceus* | AKXB | NA |
| Bryophyta | Hornworts | Dendrocerotaceae | *Nothoceros aenigmaticus* | DXOU | NA |
| Bryophyta | Hornworts | Dendrocerotaceae | *Nothoceros vincentianus* | TCBC | NA |
| Bryophyta | Hornworts | Dendrocerotaceae | *Megaceros flagellaris* | UCRN | NA |
| Bryophyta | Hornworts | Leiosporocerotales | *Leiosporoceros dussii* | ANON | NA |
| Bryophyta | Hornworts | Notothyladales | *Phaeoceros carolinianus* | RXRQ | NA |
| Bryophyta | Hornworts | Notothyladales | *Phaeoceros carolinianus* | WCZB | NA |
| Bryophyta | Hornworts | Notothyladales | *Phaeoceros carolinianus* | WEEQ | NA |
| Bryophyta | Liverworts | Blasiaceae | *Blasia sp.* | AEXY | NA |
| Bryophyta | Liverworts | Calypogeiaceae | *Calypogeia fissa* | RTMU | NA |
| Bryophyta | Liverworts | Frullaniaceae | *Frullania sp.* | TGKW | NA |
| Bryophyta | Liverworts | Schistochilaceae | *Schistochila sp.* | LGOW | NA |
| Bryophyta | Liverworts | Scapaniaceae | *Barbilophozia barbata* | OFTV | NA |
| Bryophyta | Liverworts | Scapaniaceae | *Scapania nemorosa* | IRBN | NA |
| Bryophyta | Liverworts | Plagiochilaceae/Fissidentaceae | *mixed-species* | NWQC | NA |
| Bryophyta | Liverworts | Frullaniaceae | *Frullania spp.* | CHJJ | NA |
| Bryophyta | Liverworts | Ptilidiaceae | *Ptilidium pulcherrimum* | HPXA | NA |
| Bryophyta | Liverworts | Porellaceae | *Porella pinnata* | UUHD | NA |
| Bryophyta | Liverworts | Marchantiaceae | *Marchantia paleacea* | HMHL | NA |
| Bryophyta | Liverworts | Marchantiaceae | *Marchantia paleacea* | IHWO | NA |
| Bryophyta | Liverworts | Marchantiaceae | *Marchantia polymorpha* | JPYU | NA |
| Bryophyta | Liverworts | Marchantiaceae | *Marchantia emarginata* | TFYI | NA |
| Bryophyta | Liverworts | Conocephalaceae | *Conocephalum conicum* | ILBQ | NA |
| Bryophyta | Liverworts | Monocleaceae | *Monoclea gottschei* | TFDQ | NA |
| Bryophyta | Liverworts | Lunulariaceae | *Lunularia cruciata* | TXVB | NA |
| Bryophyta | Liverworts | Ricciaceae | *Ricciocarpos natans* | WJLO | NA |
| Bryophyta | Liverworts | Metzgeriaceae | *Metzgeria crassipilis* | NRWZ | NA |
| Bryophyta | Liverworts | Pallaviciniaceae | *Pallavicinia lyellii* | YFGP | NA |
| Bryophyta | Liverworts | Pelliaceae | *Pellia neesiana* | JHFI | NA |
| Bryophyta | Liverworts | Pelliaceae | *Pellia cf. epiphylla* | PIUF | NA |
| Bryophyta | Liverworts | Pelliaceae | *Noteroclada confluens* | YPSN | NA |
| Bryophyta | Liverworts | Radulaceae | *Radula lindenbergiana* | BNCU | NA |
| Bryophyta | Liverworts | Porellaceae | *Porella navicularis* | KRUQ | NA |
| Bryophyta | Liverworts | Sphaerocarpaceae | *Sphaerocarpos texanus* | HERT | NA |
| Bryophyta | Liverworts | Treubiaceae | *Treubia lacunosa* | FITN | NA |
| Bryophyta | Mosses | Andreaeaceae | *Andreaea rupestris* | WOGB | NA |
| Bryophyta | Mosses | Bartramiaceae | *Philonotis fontana* | ORKS | NA |
| Bryophyta | Mosses | Aulacomniaceae | *Aulacomnium heterostichum* | WNGH | NA |
| Bryophyta | Mosses | Mielichhoferiaceae | *Plagiomnium insigne* | BGXB | NA |
| Bryophyta | Mosses | Bryaceae | *Bryum argenteum* | JMXW | 1 |
| Bryophyta | Mosses | Buxbaumiaceae | *Buxbaumia aphylla* | HRWG | NA |
| Bryophyta | Mosses | Ditrichaceae | *Ceratodon purpureus* | FFPD | NA |
| Bryophyta | Mosses | Dicranaceae | *Dicranum scoparium* | NGTD | NA |
| Bryophyta | Mosses | Leucobryaceae | *Leucobryum albidum* | VMXJ | NA |
| Bryophyta | Mosses | Leucobryaceae | *Leucobryum glaucum* | RGKI | NA |
| Bryophyta | Mosses | Diphysciaceae | *Diphyscium foliosum* | AWOI | NA |
| Bryophyta | Mosses | Encalyptaceae | *Encalypta streptocarpa* | KEFD | 1 |
| Bryophyta | Mosses | Bryaceae | *Rosulabryum cf. capillare* | XWHK | NA |
| Bryophyta | Mosses | Funariaceae | *Physcomitrium sp.* | YEPO | NA |
| Bryophyta | Mosses | Scouleriaceae | *Scouleria aquatica* | BPSG | NA |
| Bryophyta | Mosses | Grimmiaceae | *Racomitrium varium* | RDOO | NA |
| Bryophyta | Mosses | Hedwigiaceae | *Hedwigia ciliata* | YWNF | 1 |
| Bryophyta | Mosses | Calliergonaceae | *Calliergon cordifolium* | TAVP | 1 |
| Bryophyta | Mosses | Leucodontaceae | *Leucodon julaceus* | IGUH | NA |
| Bryophyta | Mosses | Leucodontaceae | *Leucodon brachypus* | ZACW | NA |
| Bryophyta | Mosses | Brachytheciaceae | *Rhynchostegium serrulatum* | JADL | NA |

| | | | | | |
|---|---|---|---|---|---|
| Bryophyta | Mosses | Pylaisiaceae | *Stereodon subimponens* | LNSF | NA |
| Bryophyta | Mosses | Neckeraceae | *Neckera douglasii* | TMAJ | 1 |
| Bryophyta | Mosses | Hylocomiaceae | *Loeskeobryum brevirostre* | WSPM | NA |
| Bryophyta | Mosses | Climaciaceae | *Climacium dendroides* | MIRS | NA |
| Bryophyta | Mosses | Fontinalaceae | *Fontinalis antipyretica* | DHWX | 1 |
| Bryophyta | Mosses | Myuriaceae | *Anomodon attenuatus* | QMWB | 1 |
| Bryophyta | Mosses | Brachytheciaceae | *Claopodium rostratum* | VBMM | NA |
| Bryophyta | Mosses | Orthotrichaceae | *Orthotrichum lyellii* | CMEQ | 1 |
| Bryophyta | Mosses | Polytrichaceae | *Polytrichum commune* | SZYG | NA |
| Bryophyta | Mosses | Polytrichaceae | *Atrichum angustatum* | ZTHV | NA |
| Bryophyta | Mosses | Pottiaceae | *Syntrichia princeps* | GRKU | NA |
| Bryophyta | Mosses | Sphagnaceae | *Sphagnum lescurii* | GOWD | NA |
| Bryophyta | Mosses | Sphagnaceae | *Sphagnum palustre* | RCBT | NA |
| Bryophyta | Mosses | Sphagnaceae | *Sphagnum recurvum* | UHLI | NA |
| Bryophyta | Mosses | Takakiaceae | *Takakia lepidozioides* | SKQD | NA |
| Bryophyta | Mosses | Tetraphidaceae | *Tetraphis pellucida* | HVBQ | NA |
| Bryophyta | Mosses | Timmiaceae | *Timmia austriaca* | ZQRI | NA |
| Lycophyta | Isoetales | Isoetaceae | *Isoetes sp.* | PYHZ | NA |
| Lycophyta | Isoetales | Isoetaceae | *Isoetes tegetiformans* | PKOX | NA |
| Lycophyta | Lycopodiales | Huperziaceae | *Huperzia myrsinites* | CBAE | NA |
| Lycophyta | Lycopodiales | Huperziaceae | *Phylloglossum drummondii* | ZZEI | NA |
| Lycophyta | Lycopodiales | Lycopodiaceae | *Lycopodium annotinum* | ENQF | NA |
| Lycophyta | Lycopodiales | Lycopodiaceae | *Huperzia squarrosa* | GAON | NA |
| Lycophyta | Lycopodiales | Lycopodiaceae | *Huperzia lucidula* | GKAG | NA |
| Lycophyta | Lycopodiales | Lycopodiaceae | *Huperzia selago* | GTUO | NA |
| Lycophyta | Lycopodiales | Lycopodiaceae | *Huperzia selago* | NYBX | NA |
| Lycophyta | Lycopodiales | Lycopodiaceae | *Lycopodium deuterodensum* | PQTO | NA |
| Lycophyta | Lycopodiales | Lycopodiaceae | *Lycopodiella appressa* | ULKT | NA |
| Lycophyta | Lycopodiales | Lycopodiaceae | *Pseudolycopodiella caroliniana* | UPMJ | NA |
| Lycophyta | Lycopodiales | Lycopodiaceae | *Dendrolycopodium obscurum* | XNXF | NA |
| Lycophyta | Lycopodiales | Lycopodiaceae | *Diphasiastrum digitatum* | WAFT | NA |
| Lycophyta | Selaginellales | Selaginellaceae | *Selaginella lepidophylla* | ABIJ | NA |
| Lycophyta | Selaginellales | Selaginellaceae | *Selaginella wallacei* | JKAA | NA |
| Lycophyta | Selaginellales | Selaginellaceae | *Selaginella willdenowii* | KJYC | NA |
| Lycophyta | Selaginellales | Selaginellaceae | *Selaginella selaginoides* | KUXM | NA |
| Lycophyta | Selaginellales | Selaginellaceae | *Selaginella apoda* | LGDQ | NA |
| Lycophyta | Selaginellales | Selaginellaceae | *Selaginella kraussiana* | ZFGK | NA |
| Lycophyta | Selaginellales | Selaginellaceae | *Selaginella acanthonota* | ZYCD | NA |
| Lycophyta | Selaginellales | Selaginellaceae | *Selaginella stauntoniana* | ZZOL | NA |

*Group* Major plant lineage.
*Subgroup* Subdivision within the group.
*Family* Taxonomic family of the species.
*Species* Scientific name of the species sampled.
*Code (OneKP)* Four-letter code of the species/proteome from the OneKP database.
*# Homologues* Number of homologs identified in this study (NA = not detected).

**Table S6.** NAS domain characterization and interspecies comparison.

| *Species* | *Group* | *Id* | *Length* | *Signature DB* | *Signature Acc* | *Signature Desc* | *Start* | *End* | *e-value* | *InterPro Acc* | *InterPro Desc* |
|---|---|---|---|---|---|---|---|---|---|---|---|
| *Physcomitrella patens* | Bryopsyda | Physco_Pp3c8_970V3.1.p | 321 | Pfam | PF03059 | NAS | 39 | 315 | 1.6E-70 | IPR004298 | Nicotianamine synthase |
| *Wilcoxina mikolae* | Dikarya | KAF8247961.1 | 193 | Pfam | PF03059 | NAS | 6 | 185 | 2.7E-44 | IPR004298 | Nicotianamine synthase |
| *Vezdaea aestivalis* | Dikarya | KAI9889147.1 | 312 | Pfam | PF03059 | NAS | 27 | 309 | 4.8E-58 | IPR004298 | Nicotianamine synthase |
| *Ceratodon purpureus* | Bryopsyda | CepurGG1.10G149600.1.p | 321 | Pfam | PF03059 | NAS | 41 | 318 | 6.8E-70 | IPR004298 | Nicotianamine synthase |
| *Ginkgo biloba* | Gymnosperms | gbi_evm.model.chr9.1997 | 296 | Pfam | PF03059 | NAS | 26 | 293 | 1.4E-103 | IPR004298 | Nicotianamine synthase |
| *Oryza sativa* | Monocot (Poaceae) | OsNAS1_LOC_Os03g19427.1 | 332 | Pfam | PF03059 | NAS | 5 | 280 | 3.9E-128 | IPR004298 | Nicotianamine synthase |
| *Oryza sativa* | Monocot (Poaceae) | OsNAS3_LOC_Os07g48980.1 | 343 | Pfam | PF03059 | NAS | 21 | 294 | 1.5E-123 | IPR004298 | Nicotianamine synthase |
| *Arabidopsis thaliana* | Eudicot | AtNAS1_AT5G04950.1 | 320 | Pfam | PF03059 | NAS | 4 | 276 | 9.1E-126 | IPR004298 | Nicotianamine synthase |
| *Arabidopsis thaliana* | Eudicot | AtNAS3_AT1G09240.1 | 320 | Pfam | PF03059 | NAS | 5 | 278 | 4.8E-130 | IPR004298 | Nicotianamine synthase |
| *Zalerion maritima* | Dikarya | KAJ2897573.1 | 327 | Pfam | PF03059 | NAS | 160 | 321 | 2.6E-60 | IPR004298 | Nicotianamine synthase |
| *Kalaharituber pfeilii* | Dikarya | KAF8455802.1 | 351 | Pfam | PF03059 | NAS | 41 | 344 | 9.8E-56 | IPR004298 | Nicotianamine synthase |
| *Azolla filiculoides* | Seedless Tracheophytes | Azfi_s0005.g082023 | 327 | Pfam | PF03059 | NAS | 49 | 315 | 1.6E-80 | IPR004298 | Nicotianamine synthase |

| | |
|---|---|
| *Species* | The species from which the 3D model was obtained. |
| *Group* | Higher-level group or lineage of the species. |
| *Id* | Protein identifier (query sequence ID). |
| *Length* | Protein length (aa). |
| *Signature DB* | Source database of the match. |
| *Signature Acc* | Accession of the signature. |
| *Signature Desc* | Description of the signature (domain/family). |
| *Start* | Start position of the domain in the protein. |
| *End* | End position of the domain in the protein. |
| *e-value* | Match significance (expect value). |
| *InterPro Acc* | InterPro accession. |
| *InterPro Desc* | InterPro description (protein family/function). |

**Table S7.** Evolutionary amino acid substitution models applied in IQ-TREE analyses.

| *Family/Analysis* | *Model selected* | *Selection criterion* |
|---|---|---|
| OPT - Archaeplastida | LG+R10 | AIC |
| OPT - HGT | Q.pfam+R10 | BIC |
| NAS - Archaeplastida | JTT+F+R6 | AIC |
| NAS - HGT | JTT+F+R10 | AIC |

| | |
|---|---|
| *Family/Analysis* | Proetin family or type of analysis. |
| *Model* | Best-fitting substitution model chosen for the alignment. |
| *Selection criterion* | Information criterion used to select the model. |

**Table S8.** Structural trimming coordinates for three-dimensional OPT models.

| *Species* | *Group* | *ID* | *Clade* | *N-terminal region (start)* | *N-terminal region (end)* | *Middle region (start)* | *Middle region (end)* | *Cut length (aa)* |
|---|---|---|---|---|---|---|---|---|
| *Hordeum vulgare* | Monocot (Poaceae) | HvYSL1 - 7WSR (Cryo-EM Model) | YSL | 1 | 45 | 360 | 392 | 32 |
| *Hordeum vulgare* | Monocot (Poaceae) | HvYSL1 (AlphaFold Model) | YSL | 1 | 45 | 360 | 392 | 32 |
| *Klebsormidium nitens* | Charophyta | kfl00583_0020 | PT | 1 | 129 | 478 | 493 | 15 |
| *Ginkgo biloba* | Gymnosperms | gbi_evm.model.chr9.489 | PT | 1 | 45 | 400 | 418 | 18 |
| *Oryza sativa* | Monocot (Poaceae) | LOC_Os03g54000.1 | PT | 1 | 45 | 417 | 435 | 18 |
| *Brachypodium dystachion* | Monocot (Poaceae) | Bradi1g08421.1.p | PT | 1 | 45 | 406 | 425 | 19 |
| *Sorghum bicolor* | Monocot (Poaceae) | Sobic.001G087700.1.p | PT | 1 | 45 | 406 | 425 | 19 |
| *Arabidopsis thaliana* | Eudicot | AT4G16370.1 | PT | 1 | 39 | 396 | 415 | 20 |
| *Vitis vinifera* | Eudicot | VIT_200s0259g00120.1 | PT | 1 | 45 | 409 | 425 | 16 |
| *Solanum lycopersicum* | Eudicot | Solyc11g012700.1.1 | PT | 1 | 54 | 414 | 432 | 18 |
| *Gongronella butleri* | Mucoromycota | KAI8064712.1 | PT | 1 | 69 | 416 | 432 | 16 |
| *Dunaliella salina* | Chlorophyta | Dunsal1_6490_Dusal.0238s00016.1 / KAF5836148.1 | YSL | 1 | 41 | 323 | 425 | 102 |
| *Ginkgo biloba* | Gymnosperms | gbi_evm.model.chr6.4 | YSL | 1 | 138 | 453 | 491 | 38 |
| *Oryza sativa* | Monocot (Poaceae) | LOC_Os02g43410.1 | YSL | 1 | 41 | 354 | 386 | 32 |
| *Arabidopsis thaliana* | Eudicot | AT4G24120.1 | YSL | 1 | 43 | 358 | 388 | 30 |
| *Vitis vinifera* | Eudicot | VIT_202s0025g02500.1 | YSL | 1 | 29 | 345 | 376 | 31 |
| *Solanum lycopersicum* | Eudicot | Solyc08g083060.2.1 | YSL | 1 | 29 | 346 | 378 | 32 |
| *Azolla filiculoides* | Seedless Tracheophytes | Azfi_s0001.g000194 | PT | 1 | 53 | 390 | 407 | 17 |
| *Azolla filiculoides* | Seedless Tracheophytes | Azfi_s0665.g081749 | YSL | 1 | 127 | 451 | 481 | 30 |
| *Physcomitrella patens* | Bryophyta | Physco_Pp3c10_15470V3.1.p | PT | 1 | 49 | 406 | 422 | 16 |
| *Physcomitrella patens* | Bryophyta | Physco_Pp3c7_20710V3.1.p | YSL | 1 | 75 | 397 | 427 | 30 |
| *Brachypodium dystachion* | Monocot (Poaceae) | Bradi5g17220.2.p | YSL | 1 | 33 | 350 | 378 | 28 |
| *Sorghum bicolor* | Monocot (Poaceae) | Sobic.006G164300.1.p | YSL | 1 | 78 | 398 | 425 | 27 |
| *Dispira simplex* | Zoopagomycota | KAJ1661025.1 | PT | 1 | 55 | 444 | 461 | 17 |
| *Achlya hypogyna* | SAR | OQR94115.1 | YSL | 1 | 42 | 348 | 377 | 29 |
| *Emiliania huxleyi* | Haptophyta | XP_005793885.1 | YSL | NA | NA | 309 | 357 | 48 |
| *Planctomycetota bacterium* | Bacteria | HZN60398.1 | YSL | 1 | 17 | NA | NA | NA |

| | |
|---|---|
| *Species* | The species from which the 3D model was obtained. |
| *Group* | Higher-level group or lineage of the species. |
| *ID* | Identifier of the protein sequence or structural model. |
| *Clade* | Functional or evolutionary clade assignment of the protein. |
| *N-terminal region (start)* | Start amino acid position of the N-terminal segment. |
| *N-terminal region (end)* | End amino acid position of the N-terminal segment. |
| *Middle region (start)* | Start amino acid position of the middle segment. |
| *Middle region (end)* | End amino acid position of the middle segment. |
| *Cut length (aa)* | Number of amino acids removed or considered as a cut length in the analysis. |
